# Natural selection acting on complex traits hampers the predictive accuracy of polygenic scores in ancient samples

**DOI:** 10.1101/2024.09.10.612181

**Authors:** Valeria Añorve-Garibay, Emilia Huerta-Sanchez, Mashaal Sohail, Diego Ortega-Del Vecchyo

## Abstract

The prediction of phenotypes from ancient humans has gained interest due to its potential to investigate the evolution of complex traits. These predictions are commonly performed using polygenic scores computed with DNA information from ancient humans along with genome-wide association studies (GWAS) data from present-day humans. However, numerous evolutionary processes could impact the prediction of phenotypes from ancient humans based on polygenic scores. In this work we investigate how natural selection impacts phenotypic predictions on ancient individuals using polygenic scores. We use simulations of an additive trait to analyze how natural selection impacts phenotypic predictions with polygenic scores. We simulate a trait evolving under neutrality, stabilizing selection and directional selection. We find that stabilizing and directional selection have contrasting effects on ancient phenotypic predictions. Stabilizing selection accelerates the loss of large-effect alleles contributing to trait variation. Conversely, directional selection accelerates the loss of small and large-effect alleles that drive individuals farther away from the optimal phenotypic value. These effects result in specific shared genetic variation patterns between ancient and modern populations which hamper the accuracy of polygenic scores to predict phenotypes. Furthermore, we conducted simulations that include realistic strengths of stabilizing selection and heritability estimates to show how natural selection could impact the predictive accuracy of ancient polygenic scores for two widely studied traits: height and body mass index. We emphasize the importance of considering how natural selection can decrease the reliability of ancient polygenic scores to perform phenotypic predictions on an ancient population.

## Introduction

Genome-wide association studies (GWAS) with large sample sizes and deep phenotyping have identified thousands of loci associated with complex traits and diseases ^1–4^. These associations have enabled the possibility of computing polygenic scores (PS), which represent the genetic contribution of single nucleotide polymorphisms (SNPs) to a heritable trait. Researchers have used polygenic scores as a tool to develop predictive models for inferring phenotypes and assessing individuals’ genetic risk to exhibit different phenotypic conditions^5^. The ability to sequence genomes from past human remains has allowed the analysis of ancient genotypes using polygenic scores to predict phenotypes that cannot be observed directly^6^. These polygenic score analyses have been conducted using allele effect sizes estimated with genomic data from present-day cohorts. The ability of polygenic scores to predict ancient phenotypes using ancient DNA extracted from human tissues is an area of recent interest due to its potential to investigate the evolution of anthropometric measurements such as height^7–9^ and to analyze the temporal prevalence of different diseases and conditions^10^.

Current research has demonstrated that polygenic scores can result in poor predictions among contemporary individuals. Some of the factors compromising the predictive power of polygenic scores include population stratification, changes in allele frequencies between populations, and environmental heterogeneity^11–15^. However, few studies have evaluated the impact of evolutionary factors such as natural selection on the predictive accuracy of polygenic scores on both contemporary and ancient individuals^16–18^. A previous study has shown that stabilizing selection is acting on 26 out of 70 traits analyzed in both sexes from the UK Biobank data^19^ and a more recent study showed that stabilizing selection acts on 21 out of 27 analyzed traits^20^. These results suggest that this evolutionary force needs to be considered when performing analysis of ancient phenotypic predictions due to its action on many complex traits. Previous work demonstrated that stabilizing selection reduces the predictive accuracy of polygenic scores in present-day populations not represented in GWAS samples^18^. However, to our knowledge there has not been any research analyzing how stabilizing selection impacts the predictive accuracy of polygenic scores for ancient individuals. On the other hand, previous work evaluated how directional selection impacts the predictive accuracy of ancient traits^17^ but we currently lack an understanding of differences on the action of stabilizing and directional selection on the predictive accuracy of polygenic scores in ancient humans.

In this work we investigate how stabilizing and directional selection impact the predictive accuracy of ancient polygenic scores when the scores are computed using ancient genotypes along with effect size estimates from a present-day population. We use forward in time simulations to model a single trait evolving under stabilizing selection or directional selection. We show that stabilizing and directional selection reduce the predictive accuracy of ancient polygenic scores even with perfectly estimated effect sizes at the causal loci of complex traits. Stabilizing selection causes the loss of high effect alleles while directional selection causes the loss of alleles that move phenotypes farther away from the phenotypic optimum. We observe a lower phenotypic predictive accuracy when the strength of stabilizing selection increases. We also find that the distribution of allele effects has an impact on the predictive accuracy of phenotypes when the traits evolve under directional selection. Moreover, we perform simulations to show how natural selection could impact the phenotypic prediction of height and body mass index in the past with polygenic scores. We find that stabilizing selection and directional selection negatively impact polygenic score accuracy despite having a simple demographic model, complete genotype data and perfect estimates of effect sizes in causal mutations. We argue that considering the impact of natural selection acting on a trait is important to avoid substantial biases in the prediction of complex traits from the past with the use of polygenic scores.

## Materials and methods

### Simulation details

We used *SLiM 4*.*1*^21^ to simulate a polygenic trait evolving under neutral evolution, stabilizing selection and directional selection. We simulated a single population with a constant population size of *N* = 10 000 diploid individuals. We simulated 20 independent regions of 25 000 bp to mimic the human nuclear gene median length^22^. We set the mutation and recombination rate at a value of 1 *e*^-8^ per base pair. Each independent region comprises quantitative trait loci (QTLs) where the effect size of a new allele is drawn from a normal distribution with mean *μ* = 0 and standard deviation *σ* = 0.25. We assume that the effect sizes of alleles in QTLs are additive. We defined an individual *j* true phenotype *Y*_*j*_ as *Y*_*j*_ = *G*_*j*_ + *∈*. Here *G*_*j*_ is the additive genetic value which is the sum of the additive effects of the derived alleles possessed by an individual *j* and can be estimated as 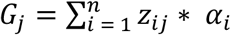 where *a*_*i*_ is the additive effect of the derived allele at SNP *i, z*_*ij*_ is the number of copies of the derived allele that the individual *j* carries at the SNP *i* and *n* is the number of new mutations. On the other hand, *ε* is an environmental variable drawn from a Gaussian random distribution with mean *μ* = 0 and standard deviation 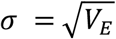.We defined the environmental variance *V*_*E*_ as *V*_*E*_ = (*V*_*G*_ − *h*^2^*V*_*G*_)/*h*^2^. Here we defined *V*_*G*_ as the genetic variance which is calculated as the variance of all the additive genetic values *G*_*j*_. We evaluated two different heritability values, *h*^2^ = {0.5, 1.0} for each evolutionary scenario.

We simulated a polygenic trait evolving under 1) neutrality, 2) stabilizing selection and 3) directional selection. In the case of neutrality, we assume that every new allele does not have an effect in the fitness of an individual. On the other hand, we modeled a scenario of stabilizing selection using a Gaussian stabilizing selection fitness function to define the fitness of an individual as

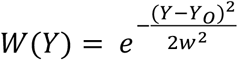

Where *Y* is the phenotype of an individual, *Y*_0_ is the optimal phenotype and *w* determines the width of the fitness peak, i.e. the strength of stabilizing selection. Larger values of *w* indicate a weaker strength of stabilizing selection^23,24^. We ran simulations of a trait evolving under stabilizing selection around a constant optimal trait value *Y*_0_ = 0 at five different strengths of selection, *w* = {1,2,3,4, 5} to represent cases going from stronger to weaker stabilizing selection, respectively. The values of *w* are in the same order of magnitude of values of *w* estimated on traits under stabilizing selection in humans^18^.

We forced a positive shift in the optimal trait value to simulate a trait evolving under directional selection. In our simulations, the population evolved under stabilizing selection with *w* = 1 and an optimal trait value of *Y*_1_ = 0 during a burn-in period of 10*N* generations. The optimal trait value then shifts to *Y*_1_ = 1 for the remainder 400 generations that we simulated. We changed the standard deviation of the QTL mutations effect sizes distribution *QTL* ∼ *N*(*μ* = 0, *σ* = {0.25, 0.025, 0.0025}) under the scenario of directional selection. For this selection scenario we computed the phenotypic mean and phenotypic variance for every generation since the optimal value shift until the present.

In all our simulations we used a burn in period of 100 000 (10*N*) generations. We randomly sampled 100 individuals from the population every 100 generations for 400 generations after the burn in period. We defined ancient sampling times as τ = 0, 100, 200, 300, 400 generations in the past. These τ values were chosen to mimic sampling times of 0, 2 900, 5 800, 8 700 and 11 600 years before the present assuming a generation time of 29 years per generation^25^. These sampling times contain the timeframe of 0 − 10 000 years before the present where the majority of the recovered genetic samples from ancient humans have been collected^26^. We record the genotype and allele QTL effect sizes of each individual we sampled at different times τ. We simulated 100 replicates for each combination of parameter values in each scenario.

We compute the *true phenotype, Y*_*j*_(τ = *x*), for each sampled ancient individual *j* at time τ = *x* as

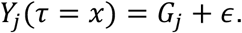

We define the *ancient polygenic scores* 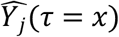 for each *j* sampled individual at time τ = *x* as

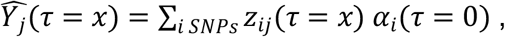

where *z*_*ij*_(τ = *x*) ∈ {0,1,2} is the number of copies of the derived allele at the *i*th SNP of the *j*th individual sampled at time τ = *x*. GWAS can only estimate the effect sizes of variants on segregating sites in present-day populations. Due to this, our estimations of 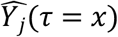 only use alleles from variants present on segregating sites in a present-day sample of 100 individuals at τ = 0. *α*_*i*_(τ = 0) is equal to 0 if the derived allele is not present in a segregating site on a present-day sample of 100 individuals. On the other hand, *α*_*i*_(τ = 0) is equal to the effect size of the variant if the derived allele is present in a segregating site on a present-day sample of 100 individuals. In our modeling framework we assume that we know the effect sizes of variants in segregating sites at time τ = 0 in a present-day sample of 100 individuals. Therefore, we know the effect sizes of all variants with a frequency equal or bigger than 0.5% on segregating sites in a present-day sample of 100 individuals. **Figure 1** summarizes the modeling framework.

**Figure 1.**
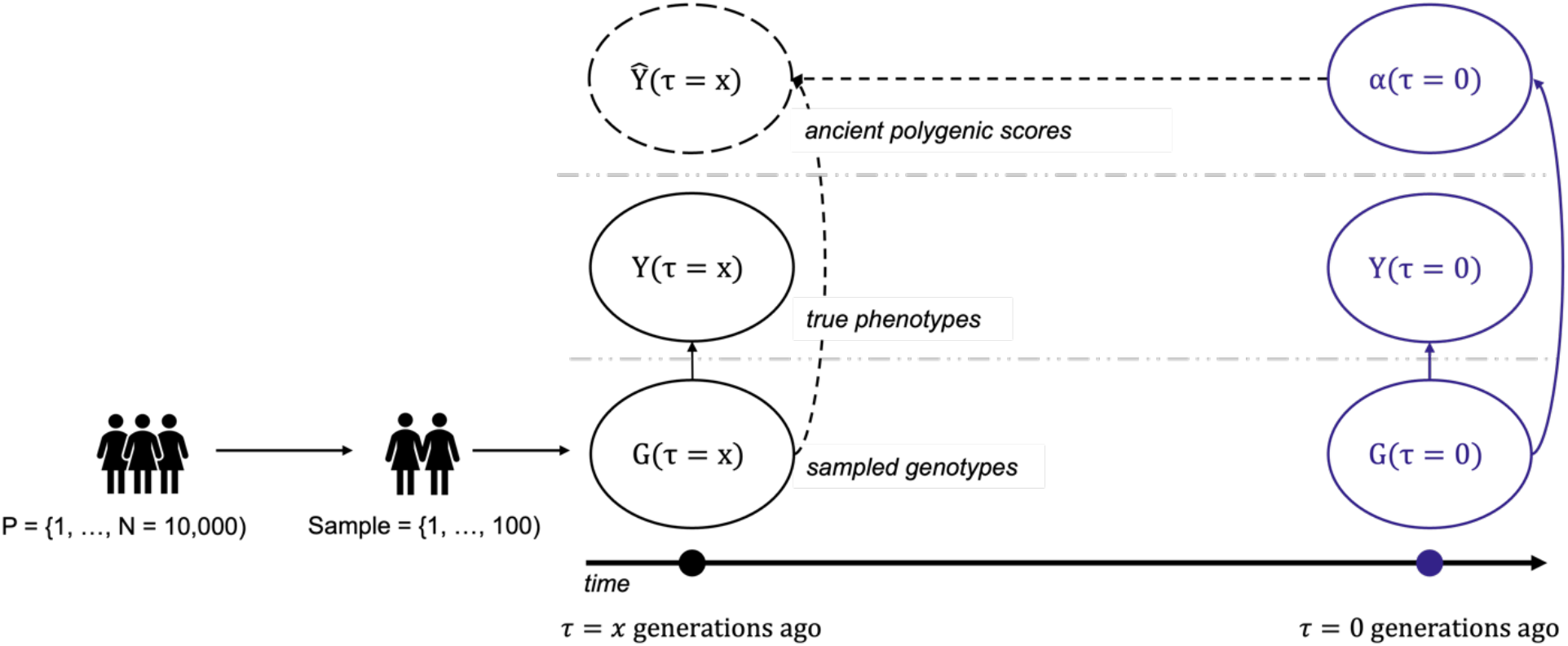
Our modeling framework to compute ancient polygenic scores (*aPS*). 100 individuals are sampled at a time point τ from the same population where a GWAS was conducted at time τ = 0 (present day). Ancient polygenic scores are computed for each individual at time τ using both their genotype data at time τ and the effect sizes *α*(τ = 0) from the present-day GWAS. Purple and solid circles represent observed values at time τ = 0 while black and solid circles represent observed values at time τ. The dashed circle represents the estimated ancient polygenic scores at time τ. This figure is inspired by Figure 1C from ^17^.

For the neutral evolution case, we computed the transition mass functions (TMF), i.e. the probability of transitioning from a specific number of alleles at one point in time to a different number of alleles at a point in the future. We used fastDTWF^27^, which is a tool to compute likelihoods and transition probabilities under the discrete-time Wright-Fisher model. We assumed a population size of 20 000 haploids with a mutation rate of 1 *e* − 8 and set the selection coefficient to *s* = 0 to explore a scenario where the alleles evolve under neutrality. Our initial allele frequencies were based on a set ranging from 0 to 0.02 in 0.001 steps, as approximately 90% of all our simulated alleles on QTLs for the neutral case at *h*^2^ = 1.0 have population frequencies between 0 and 0.02. We computed the transition probabilities for the alleles to be lost on 400 generations.

### Accuracy metrics

We used two statistics to assess the accuracy of the ancient polygenic score in approximating the true phenotype. First, we used the coefficient of determination, *r*^2^, to measure the error of ancient polygenic scores to predict phenotypes. *r*^2^ is the squared value of the Pearson’s correlation coefficient, *r* and is defined as:

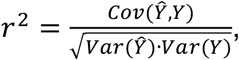

where *Y* and *Ŷ* are the *n*-vectors of true phenotypes *Y*_*j*_(τ = *x*) and ancient polygenic scores 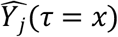 at some time τ = *x*, respectively, of all individuals *j*.

We also used another metric defined as the mean-squared error which is equal to:

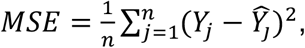

where *n* is the sample size, *Y*_*j*_ is the *j* element of the n-vector of the true phenotypes *Y*(τ = *x*) and *Y*_*j*_ is the is the *j* element of the n-vector of the ancient polygenic scores 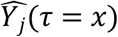 at some time τ = *x* of all individuals *j*.

## Results

### Accuracy of ancient polygenic scores on a trait evolving under neutrality

We employed a modeling framework (see **Methods**) to analyze the ability of polygenic scores to predict the true phenotype of an ancient individual sampled at a point in time τ. We used the effect sizes of QTL mutations which are assumed to be perfectly estimated from an association study (GWAS) performed in the present (τ = 0). We assume that it is only possible to estimate effect sizes from variants present on segregating sites on the GWAS. We simulated a trait evolving under neutrality (see **Methods: Simulation Details**) and we sampled individuals from five different sampling times spanning τ = {0, 100, 200, 300, 400} generations ago from the present. Remarkably, we found that ancient polygenic scores accurately predict the true phenotype of an ancient individual at the five different ancient sampling times tested when assuming a heritability value of *h*^2^ = 1.0. The values of *r*^2^(*Y*, *Ŷ*) were higher than 0.93 at τ = 400, 300, 200, 100, and 0 generations ago, respectively **(Figure 2A)**. In addition, we observed that *MSE*(*Y*, *Ŷ*) values tend to decrease linearly as we move forward in time with the largest *MSE*(*Y*, *Ŷ*) values tending to occur at τ = 400 generations ago **(Figure 2B)**. In addition, we found that ancient polygenic scores display lower *r*^2^(*Y*, *Ŷ*) values when assuming a heritability value of *h*^2^ = 0.5 compared to simulations done with a heritability value of *h*^2^ = 1.0 . We observed that *MSE*(*Y*, *Ŷ*) values are higher in simulations where *h*^2^ = 0.5 compared to simulations performed where *h*^2^ = 1.

**Figure 2.**
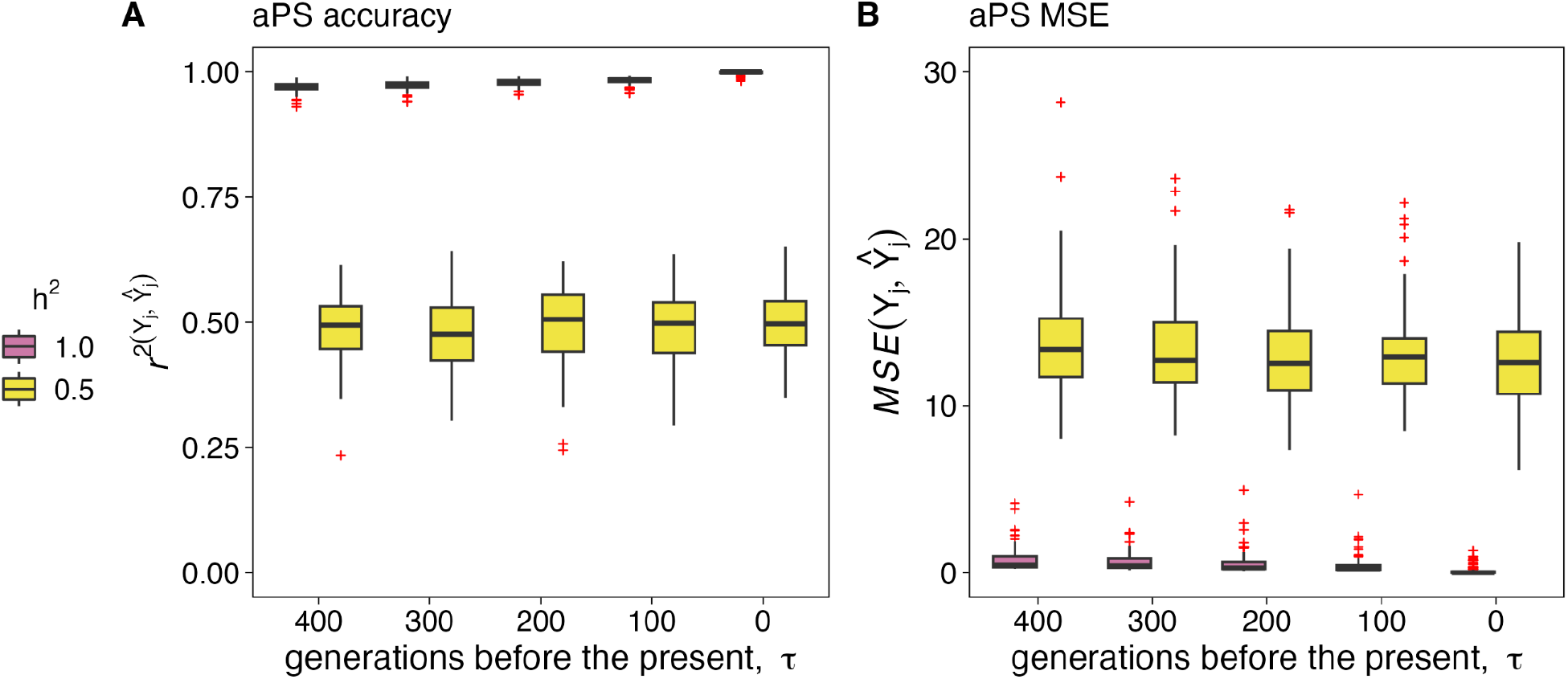
Ancient polygenic scores (aPS) accuracy (*r*^*2*^) and Mean Squared Error (MSE) for a trait evolving under neutrality. We simulated a population with a neutral trait that has a heritability value of *h*^2^ = 1.0 (pink) and *h*^2^ = 0.5 (yellow). Boxplots show the distribution of **A)** *r*^2^(*Y*, *Ŷ*), and **B)** the Mean Squared Error (MSE), *MSE*(*Y*, *Ŷ*), between the true phenotypic values and their predicted ancient polygenic scores in a sample of 100 individuals taken at five different points in times τ = 0, 100, 200, 300, 400 generations before the present. We performed 100 simulation replicates for both a heritability value *h*^2^ = 1 and *h*^2^ = 0.5 . Red crosses represent outliers.

Our model assumes that not all segregating sites will be present in both ancient and present-day sampled individuals due to genetic drift. Therefore, we evaluated the effect sizes of the QTL mutations that are conserved (i.e. mutations that remain present in both ancient and present-day sampled individuals) and lost in samples of 100 individuals taken at the earliest sampling time and the present-day sampling time (τ = 400 and 0 generations ago, respectively). We observed that the distribution of both conserved and lost alleles has a unimodal shape (**Figure S1**). We also find that the majority of QTL mutations that have a larger contribution to the trait are conserved over time in a span of 400 generations (**Figure S1**). We computed the probability of transitioning from a given allele frequency at one point in time to a different frequency at some point in the future under the forward-in-time discrete-time neutral Wright-Fisher (DTWF) model^27^ to further understand the dynamics of conserved and lost mutations. We saw that approximately 90% of the lost mutations in a span of 400 generations have population frequencies between 0 and 0.02 (**Figure S2A**). We observed that the probability of losing an allele, i.e. transitioning from *f* to *f* = 0 in τ generations increases as we move forward in time. However, as the initial frequency *f* increases, the probability of losing an allele at frequency *f* drastically decreases even at τ = 400 generations (**Figure S2B**). These results are concordant with our observation that most of the lost alleles have allele frequencies smaller than 2% (**Figure S2A**). This indicates that, under neutrality, the allele frequencies of segregating alleles must be low to be removed by genetic drift alone on the timeframe explored.

### Accuracy of ancient polygenic scores on a trait evolving under stabilizing selection

Several human complex traits evolve under stabilizing selection^19,20^. Therefore, we expanded our baseline modeling framework to simulate a trait evolving under Gaussian stabilizing selection^23,24^ to analyze whether ancient polygenic scores predictive accuracy will vary from our neutral model results. We simulated a single trait evolving under Gaussian stabilizing selection where an individual’s fitness is defined as a fitness function,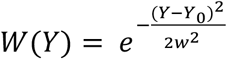 *j*, where *Y* is the individual’s phenotype, *Y*_0_ is the optimal phenotypic value and *w* is a parameter that measures the width of the fitness function and determines the strength of selection acting on phenotypes (see **Methods: Simulation details**). We assumed an evolutionary scenario in which the optimal phenotypic value, *Y*_0_ = 0, remains constant through time. We conducted simulations under different stabilizing selection strengths, *w* = {1, 2, 3, 4, 5}, ranging from stronger to weaker stabilizing selection values. We selected our *w* estimates to be on the same order of magnitude as previous estimates of *w* on real data^19^ as shown previously^18^.

We found that simulating a trait evolving under stabilizing selection at a heritability value of *h*^2^ = 1.0 decreases ancient polygenic scores predictive accuracy in contrast a trait evolving under neutrality. We observed that *r*^2^(*Y*, *Ŷ*) drops from 1 (in the present) to roughly 0.75 and 0.25 in 100 generations in the past under the different strengths of stabilizing selection that we used (**Figure 3A**). This drop continues gradually until it reaches our most ancient sampling time at 400 generations in the past. We observed that the drop in accuracy is larger under a stronger stabilizing selection force, *w* = 1, as *r*^2^(*Y*, *Ŷ*) drops from 1 (in the present) to roughly 0.27 in 100 generations in the past and to 0.21, 0.20 and 0.19 in 200, 300 and 400 generations ago, respectively (**Figure 3A**). Assuming a lower heritability of the trait, *h*^2^ = 0.5, we observed that ancient polygenic scores display poorer prediction accuracy at the five different strengths of stabilizing selection that we analyzed. We observed that ancient polygenic scores accuracy drastically decreases from a *r*^2^(*Y*, *Ŷ*) equal to 0.51 in the present to 0.13, 0.11, 0.1 and 0.08 at τ = 100, 200, 300, and 400 generations ago, respectively, when *w* = 1 and *h*^2^ = 0.5 (**Figure 3A**). In addition, we observed that *MSE*(*Y*, *Ŷ*) values appear to decrease linearly as we move forward to the present (τ = 0) at the five different *w* values. Interestingly, we find that the highest *MSE*(*Y*, *Ŷ*) values are observed at *w* = 5 and the lowest *MSE*(*Y*, *Ŷ*) values appear on simulations with *w* = 1. Therefore, we see that we have low *r*^2^(*Y*, *Ŷ*) and *MSE*(*Y*, *Ŷ*) values on traits simulated with the highest strength of stabilizing selection. This situation is a notable contrast to the results with complex traits evolving under neutrality (**Figure 2**) where cases with a low *r*^2^(*Y*, *Ŷ*) value exhibit a high *MSE*(*Y*, *Ŷ*) value and vice versa. The inspection of a simulation replicate sheds more light on this result and reveals that phenotypic values are clustered around a small number of *Y* values that are not well predicted by *Ŷ* values that do not show large variations under a strong stabilizing selection (**Figure S3**). This leads to low *MSE*(*Y*, *Ŷ*) and *r*^2^(*Y*, *Ŷ*) values when there is a high strength of stabilizing selection. Conversely, we observe high *MSE* values consistent with a wider distribution of phenotypic values under a weak stabilizing selection strength and we also observe a high *r*^2^(*Y*, *Ŷ*) (**Figure S3**). This result suggests that the magnitude of *MSE*(*Y*, *Ŷ*) depends on the strength of stabilizing selection. It also shows that different error metrics must be evaluated to define the prediction accuracy of ancient polygenic scores.

**Figure 3.**
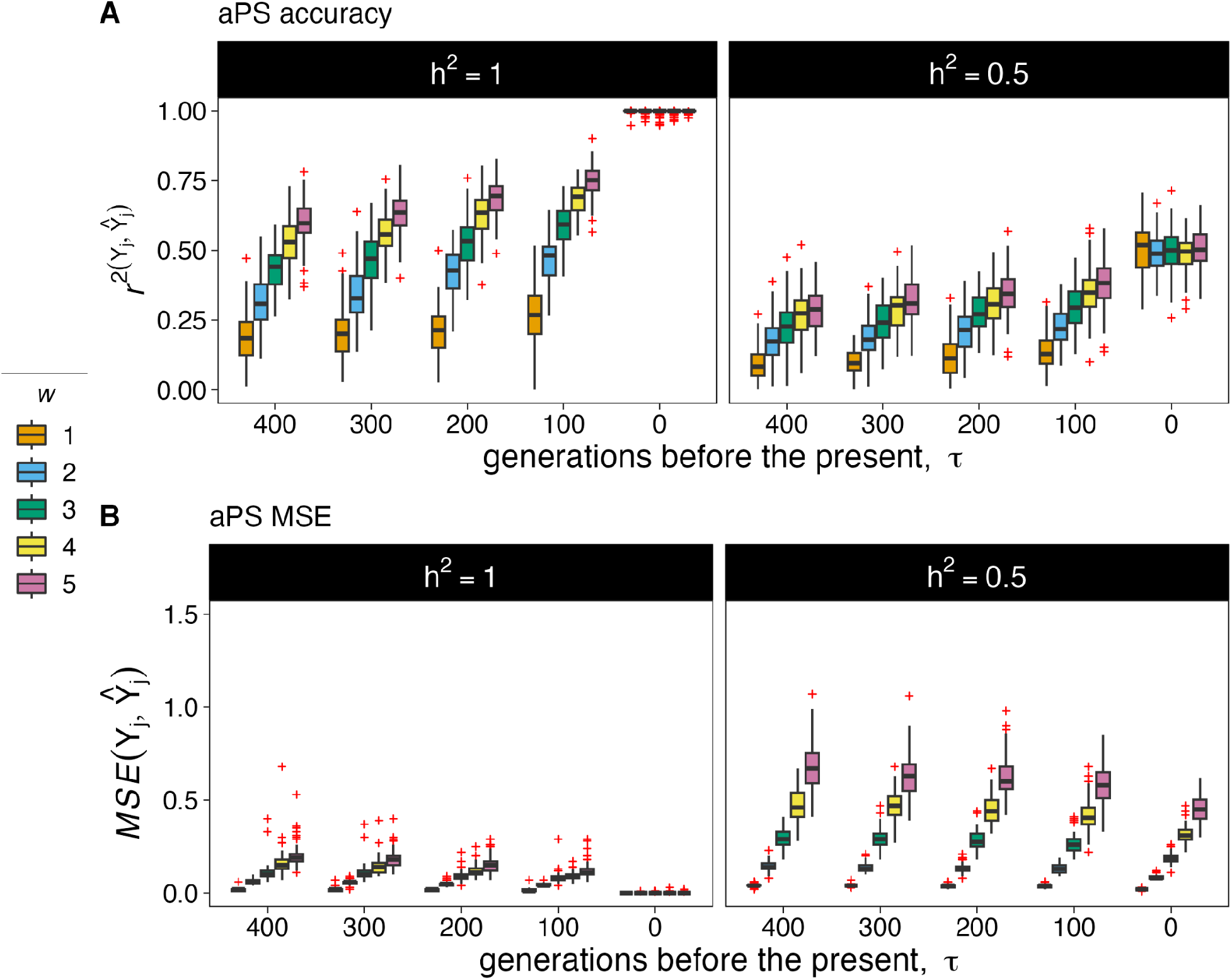
Ancient polygenic scores (aPS) accuracy (*r*^*2*^) and Mean Squared Error (MSE) for a trait evolving under stabilizing selection. We simulated a population with a trait evolving under stabilizing selection that has heritability value of *h*^2^ = 1.0 (left) and *h*^2^ = 0.5 (right). The trait is evolving under stabilizing selection at five different strengths of selection based on the parameter *w* = 1, 2, 3, 4, 5 ranging from strong to weak selection, respectively. Boxplots show the distribution of **A)** *r*^2^(*Y*, *Ŷ*), and **B)** the Mean Squared Error (MSE), *MSE*(*Y*, *Ŷ*), between the true phenotypic values and their predicted ancient polygenic scores in a sample of 100 individuals taken at different points in time, τ = 0, 100, 200, 300, 400 generations before the present in 100 simulation replicates. Simulations with a highest strength of stabilizing selection, i.e. smaller *w* values, reduce phenotypic variation and decrease *MSE*(*Y*, *Ŷ*) as seen in **Figure S3** and explained on the main text.

We then analyzed the effect sizes of the QTL mutations that are lost and conserved between the earliest sampling time and the present-day sampling time (τ = 400 and 0 generations ago, respectively). We observed that the strength of stabilizing selection causes the distribution of lost and conserved QTL allele effect sizes to be narrower when *w* is smaller and, therefore, there is a stronger stabilizing selection acting on the trait (**Figure S4**). This effect is due to the tendency to have small effect sizes in QTL mutations conserved through time. We found similar results at *h*^2^ = 0.5 for the effect sizes of the QTL mutations that are lost and conserved between the earliest sampling time and the present-day sampling time (τ = 400 and 0 generations ago, respectively) (**Figure S4)**. This result suggests that stabilizing selection increases genetic differentiation through time particularly through the loss of large-effect QTL mutations which causes a decay in ancient polygenic scores accuracy through time.

### Accuracy of ancient polygenic scores on a trait evolving under directional selection

We used our baseline Gaussian stabilizing selection fitness function to model recent directional selection as a shift where the optimum phenotypic value changed from *Y*_0_ to *Y*_0_′ in the most recent 400 generations. We studied how the accuracy of ancient polygenic scores changes due to shifts on the distribution of effect sizes that is acting on the QTL mutations. We studied this impact by reducing the standard deviation of the QTL mutations effect sizes distribution by one, two and three orders of magnitude, *QTL* ∼ *N*(*μ* = 0, *σ* = {0.25, 0.025, 0.0025}).

We first observed that the population approaches the new optimum phenotypic value within approximately 50 and 150 generations when the standard deviation of the QTL mutations effect sizes distribution is equal to 0.25 and 0.025, respectively. Conversely, the population does not approach the new optimum within the 400 generation time span at the lowest standard deviation of the distribution of effect sizes inspected, *σ* = 0.0025, (**Figure S5**). In concordance with previous research^28^, we observed that the average phenotypic variance spikes as the population approaches the new optimum value. Afterwards, the phenotypic variance is reduced (**Figure S5**). The most pronounced and severe spike occurs at *σ* = 0.25 when the population rapidly reaches the new optimum. In contrast, we do not see such a larger spike at *σ* = 0.025 and *σ* = 0.0025 with the population taking longer to approach the new optimum phenotypic value (**Figure S5**).

We then investigated how ancient polygenic scores predictive accuracy acts when the trait evolves under directional selection. We observed that, at a heritability value of *h*^2^ = 1.0, ancient polygenic scores give good predictions of the true phenotype when the QTL mutations effect sizes distribution has a lower standard deviation, *σ* = {0.025, 0.0025}. We saw that the *r*^2^(*Y*, *Ŷ*) values drop from 1 to 0.93, 0.92, 0.9 and 0.64 in samples taken 100, 200, 300 and 400 generations ago when *σ* is equal to 0.025. Additionally, the *r*^2^(*Y*, *Ŷ*) values go from 1 to 0.96, 0.95, 0.9 and 0.84 in samples taken 100, 200, 300 and 400 generations when *σ* is equal to 0.0025, respectively **(Figure 4)**. On the other hand, ancient phenotypic predictions perform poorly when *σ* = 0.25 and *r*^2^(*Y*, *Ŷ*) drops from 1 (in the present) to roughly 0.65 in 100 generations in the past and to 0.51, 0.29 and 0.29 in samples taken 200, 300 and 400 generations in the past, respectively **(Figure 4)**. On simulations done with *h*^2^ = 0.5 we observed that ancient polygenic scores display lower *r*^2^(*Y*, *Ŷ*) values compared to simulations done with *h*^2^ = 1.0 at the three *σ* values inspected. In addition, we observed *MSE*(*Y*, *Ŷ*) values decrease as we move forward to the present (τ = 0) at *σ* = 0.025 and *σ* = 0.0025. Interestingly, simulations with *σ* = 0.25 show an increase in *MSE*(*Y*, *Ŷ*) from 400 to 300 generations ago. To further understand this observation, we computed both *r*^2^(*Y*, *Ŷ*) and *MSE*(*Y*, *Ŷ*), between the true phenotype, *Y*_*j*_(τ = *x*), and the predicted ancient polygenic score, 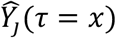, of each sampled individual *j* at times τ = 400, 390, 380, 370, 360, 350, 340, 330, 320, 310 and 300 generations before the present for simulation replicates having *σ* = 0.25 (**Figure S6**). We observed that there is a decrease in *r*^2^(*Y*, *Ŷ*) that lasts around ∼30 generations between 400 and 370 generations before the present, which is the time span the population approaches the new optimum value when *σ* = 0.25 (**Figure S5 and S6**). The values of *r*^2^(*Y*, *Ŷ*) increase as we move forward in time after 370 generations before the present.

**Figure 4.**
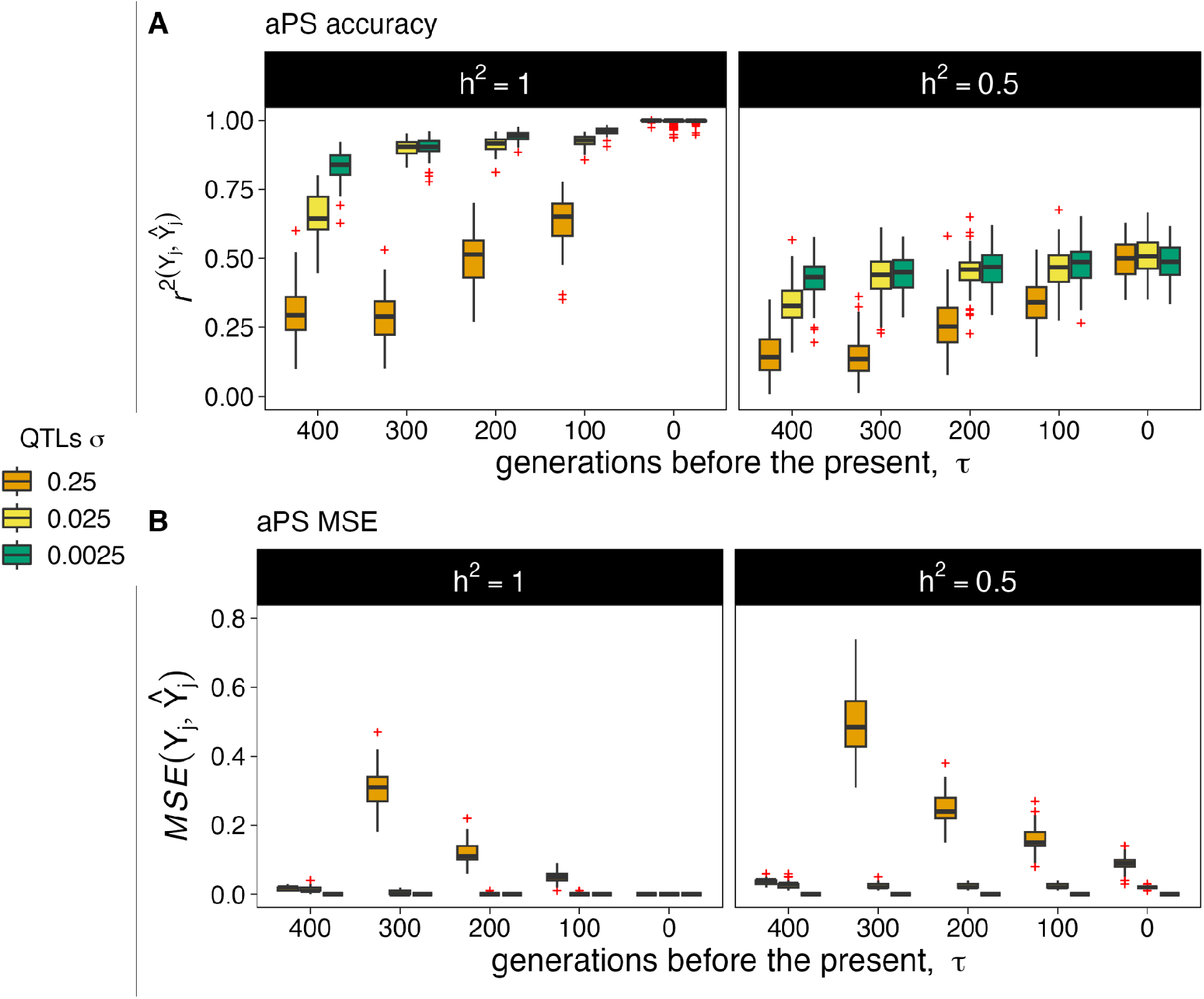
Ancient polygenic scores (aPS) accuracy (*r*^*2*^) and Mean Squared Error (MSE) for a trait evolving under directional selection. We model a trait with a heritability value of *h*^2^ = 1.0 (left) and *h*^2^ = 0.5 (right) evolving under directional selection with an optimum shift from *Y*_0_ = 0 to *Y*_0_′ = 1 over a 400 generation time span. We tested three different standard deviations of the QTL mutations effect sizes, *QTL* ∼ *N*(*μ* = 0, *σ* = {0.25, 0.025, 0.0025}), represented in orange, yellow and green, respectively. Boxplots show the distribution of **A)** *r*^2^(*Y*, *Ŷ*), and **B)** the Mean Squared Error (MSE), *MSE*(*Y*, *Ŷ*), between the true phenotypic values and their predicted ancient polygenic scores of a sample of 100 individuals at different points in times τ = 0, 100, 200, 300, 400 generations before the present in 100 simulation replicates. Red crosses represent outliers.

We analyzed the effect sizes of the QTL mutations that are lost and conserved between each ancient sampling time (τ = 400, 300, 200 and 100 generations ago, respectively) and the present-day sampling time (τ = 0 generations ago). Broadly we observed a bias where QTL mutations with negative effect sizes values tend to be more lost compared to QTL mutations with positive values which tend to be conserved (**Figure S7-S14**). Mutations with positive values move individuals closer to the new optimum value in the generation where there is a shift in the optimum phenotypic value.

### Insights into predicting the evolution of complex traits: Height and Body Mass Index (BMI)

Height and body mass index (BMI) are among the most extensively studied polygenic traits in humans and are evolving under stabilizing selection based on data from the UK Biobank^19^. We used simulations to investigate the impact of stabilizing selection on the predictive accuracy of ancient polygenic scores for these traits. Each trait evolves under stabilizing selection in our simulations based on the strength of selection given by *w* estimated from the selection gradients 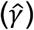 calculated previously for each trait^19^. Particularly, we used approximations to estimate *w* based on 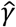 selection gradients^18^. Then we simulated height and BMI with parameters *w* = 7.28 and *w* = 6.61 and heritability values previously estimated of *h*^2^ = 0.8^29^ and *h*^2^ = 0.7^11^, respectively. In concordance with our previous results, we found that the predictive accuracy of ancient polygenic scores decreases as the time between the present-day GWAS population and the ancient population increases for both traits (**Figure 5**). This result suggests that, even with complete genotype data and perfect estimates of the effect sizes of causal mutations, polygenic scores accuracy can decay if a population is evolving under stabilizing selection.

**Figure 5.**
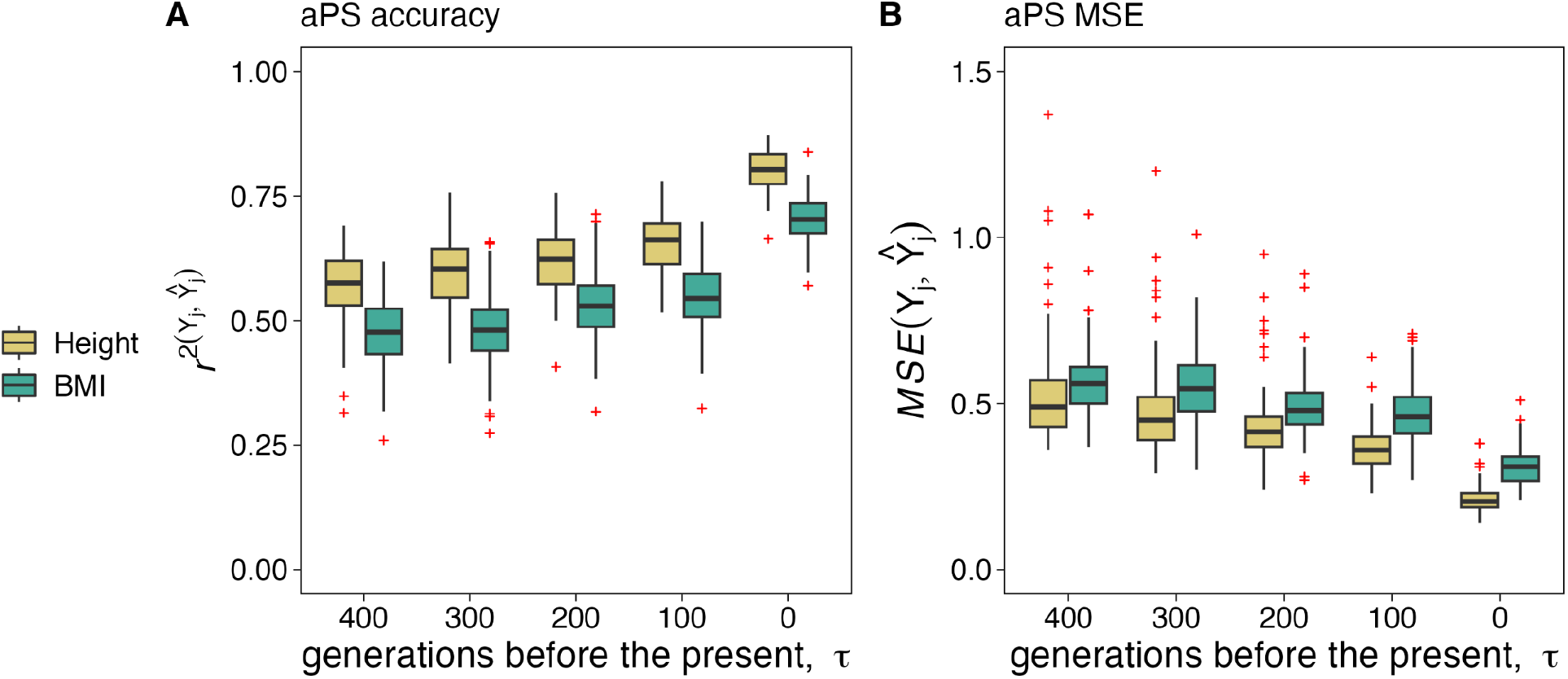
Ancient polygenic scores (aPS) accuracy (*r*^*2*^) and Mean Squared Error (MSE) for Height and BMI evolving under stabilizing selection. We simulated the evolution of height (yellow) and BMI (green). We used heritability values of *h*^2^ = 0.8 and *h*^2^ = 0.7, respectively. We determined the strength of stabilizing selection acting on each trait, *w*, based on Sanjak et al., 2017 selection gradients estimated from the UK Biobank. Boxplots show the distribution of **A)** *r*^2^(*Y*, *Ŷ*), and **B)** the Mean Squared Error (MSE), *MSE*(*Y*, *Ŷ*), between the true phenotypic values and their predicted ancient polygenic scores of a sample of 100 individuals at different points in times τ = 0, 100, 200, 300, 400 generations before the present over 100 simulation replicates. Red crosses represent outliers.

## Discussion

Inferring complex traits from genotypes in ancient samples will help to better characterize phenotypic diversity in ancient human populations. Polygenic scores provide a framework to infer ancient phenotypes. However, we still do not know all the factors that can impact the predictions of phenotypes in ancient human populations. In this work we proposed a simulation framework to investigate the impact of natural selection on the predictive accuracy of ancient polygenic scores. We show that the evolution of a phenotype under neutrality, stabilizing selection and directional selection has a different impact on the predictive accuracy of ancient polygenic scores. This reduction in the predictive accuracy is seen in samples that were taken between 0 to 400 generations ago which, assuming a generation time of 29 years^25^, contains the timeframe from 0 up to 10 000 years ago where the majority of ancient human genomes have been sampled^26^.

Our results show that we can make an accurate prediction of traits based on polygenic scores when the trait is evolving under neutrality. In our simulations we assume that we can predict the effect sizes of segregating variants with a frequency equal or larger than 0.5% frequency in the present since we take a sample of 100 individuals present-day individuals and assume that we can predict the effect sizes of all the variants segregating in this sample. It is remarkable to note we can make a very accurate prediction of neutral traits in the past knowing the effect sizes from segregating variants in that present day sample **(Figure 2)**. On the other hand, we find that stabilizing selection negatively impacts the predictive accuracy of ancient polygenic scores. This result is consistent with previous work showing that a higher strength of stabilizing selection causes more genetic differentiation among populations which negatively impacts polygenic scores accuracy in contemporary populations^18^. Similarly, we observe that stabilizing selection rapidly removes large-effect mutations within short time periods. As a result, present-day individuals become more genetically differentiated in high effect alleles from ancient individuals of the same population which is a factor that is likely to be a major contributor in the reduction of ancient polygenic score accuracy **(Figure 3)**.

Additionally, our results show that the distribution of effect sizes has an impact on the predictive accuracy of ancient phenotypic traits under directional selection. In our simulations we find that a distribution that produces a higher proportion of high effect alleles causes a higher reduction of the predictive accuracy of traits under directional selection **(Figure 4)**. This result shows that the distribution of effect sizes in the alleles acting on the trait will be important to determine the accuracy of ancient phenotypic predictions. On the other hand, we also observed that directional selection tends to preserve both small and large-effect mutations that drive individuals towards the new phenotypic optimum in the generation when there is a shift towards a new phenotypic optimum **(Figure S7-S14)**. Obtaining accurate estimates of the shape of the distribution of effect sizes will be crucial to characterize how the interaction between directional selection and the effect sizes of new mutations impacts predictions of traits on ancient individuals.

A recent paper predicted individual height variation of ancient individuals using polygenic scores^8^. They found that polygenic scores in ancient individuals can explain a modest (∼6%) but significant proportion of height variation. This finding might seem surprising considering that height is a trait with high heritability (∼80%) among present-day individuals^29^. However, as shown previously, height is evolving under stabilizing selection in individuals from the UK Biobank^19^. Here we see that stabilizing selection can reduce polygenic scores predictive accuracy based on the stabilizing selection strength estimate for height^19^ (**Figure 5**). We argue that considering stabilizing selection as a potential factor decreasing prediction accuracy will benefit the interpretation of ancient polygenic scores.

Broadly we find an interesting pattern based on the analysis of the ratio of the number of conserved QTL mutations divided by the total number of QTL mutations between the earliest sampling time and the present-day sampling time (τ = 400 and 0 generations ago, respectively) for each evolutionary scenario that we analyzed. We observed that the ratio remains constant across varying effect sizes on traits evolving under neutrality. This result shows that high effect alleles can be conserved in the time span inspected on traits evolving under neutrality (**Figure 6A**). In contrast, we showed that stabilizing selection favors the conservation of small-effect mutations while large-effect mutations are lost.

**Figure 6.**
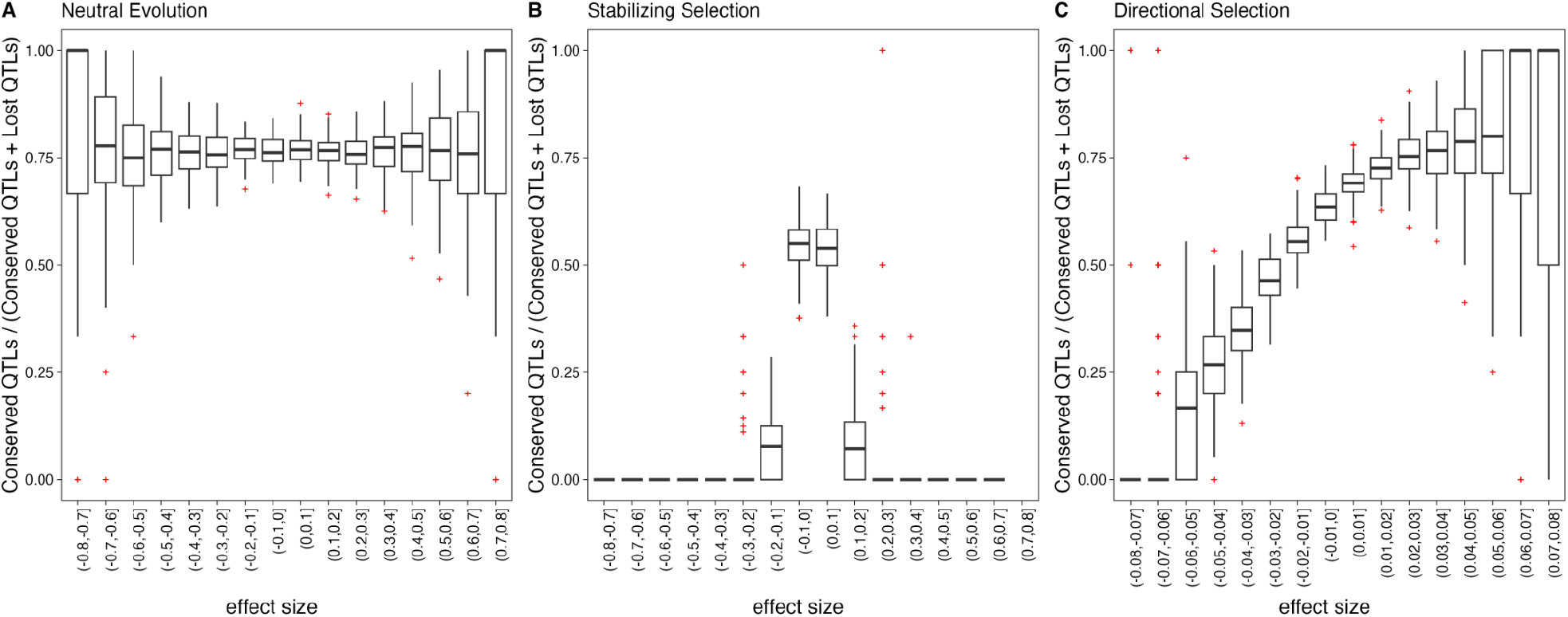
The number of conserved QTL mutations divided by the total number of QTL mutations (conserved QTLs plus lost QTLs) (Y-axis) per effect size bin (X-axis) between the earliest sampling time and the present-day sampling time (τ = 400, 0 generations ago, respectively). Results are shown for 100 replicates at heritability values of *h*^2^ = 1.0 for traits evolving under **A)** Neutrality, **B)** Stabilizing selection with a parameter *w* = 1 and **C)** Directional selection with a QTLs effect sizes distribution *QTL* ∼ *N*(*μ* = 0, *σ* = {0.025}). Red crosses represent outliers.

This pattern causes an increased genetic differentiation between ancient and present-day individuals from the same population (**Figure 6B**). Broadly, we see that in simulations done with traits evolving under stabilizing selection there is an association between a decreased accuracy of ancient polygenic scores accuracy and a higher loss of alleles with a high effect. Finally, directional selection favors the loss of negative QTL mutations. These mutations drive individuals farther away from the new phenotypic optimum in the generation when the optimum value shifts (**Figure 6C**). Therefore, natural selection changes the proportion of lost alleles based on the effect sizes of the alleles. The category of alleles lost based on their effect sizes should be considered when performing phenotypic predictions in the past since there is an association between those two factors and the type of natural selection acting on the trait.

Our simulations were done under a simple demographic model where we did not include demographic processes such as population size changes or gene flow. Here we aim to show that two evolutionary processes such as stabilizing selection and directional selection have an important impact on the prediction of ancient complex traits. Recent studies have demonstrated that stabilizing selection drives the evolution of various human complex traits^19,30,20^. On the other hand, a recent study suggests that directional selection drives the evolution of height in individuals from Sardinia^31^. Therefore, the impact of those two evolutionary processes should be considered when predicting ancient complex traits. Currently there are estimates of the impact of stabilizing selection acting on several human phenotypes^19,20^. We hope to see more studies evaluating whether complex traits exhibit signals of natural selection. It would be particularly interesting to see if the impact of natural selection acting on traits varies between population cohorts and if there are environmental factors driving the global variation in the action of natural selection.

Here we demonstrate that natural selection can hamper the predictive accuracy of ancient polygenic scores under a simple demographic model of a constant population size. Previous work has shown that the loss of alleles contributes to a decrease in the predictive accuracy of traits evolving under a neutral scenario and a directional selection scenario in a constant population size scenario^17^. Our results are consistent with that claim and, additionally, here we contrast how the predictive accuracy of ancient polygenic scores varies between neutral traits and traits that evolve under stabilizing and directional selection. Understanding the action of stabilizing selection is particularly important given its widespread effect on complex traits^19,20^. We additionally show that the action of natural selection acting on the trait impacts the alleles that tend to be lost based on their effect sizes. Finally, our simulations show that the predictive accuracy of traits can decrease on ancient samples that come from sampling times that are reflective of the period where we have more information from ancient DNA.

Finally, we acknowledge that population demographic history is an important factor impacting phenotypic variation among present-day individuals^12,14,32^ and that population demographic history coupled with natural selection can significantly impact ancient polygenic scores accuracy. We encourage future studies on ancient traits predictions to take the demographic history of each population and the impact of natural selection acting on complex traits into account. Software that can jointly model demographic history and the impact of natural selection acting on traits, e.g. SLiM^21^, should help to perform realistic simulations where the reliability of the phenotypic predictions can be quantified. We also encourage future simulation studies to include as much information as possible regarding the genetic architecture of the trait being studied. These considerations would lead us towards a more rigorous assessment of complex traits evolution.

## Acknowledgements

We thank María C. Ávila-Arcos for her valuable comments on this manuscript that helped to improve it. This work was supported by the PAPIIT-UNAM IN215524 and NIH grant R01HG012605. M.S. was supported by the PAPIIT-UNAM grant IA209024. We thank Jair S. Garcia-Sotelo, Luis Alberto Aguilar Bautista, Christian Molina-Aguilar, Alejandra Castillo, and Carina Uribe for technical assistance.

## Data availability

Code used to generate simulations, process output and plot figures can be found at: https://github.com/vagaribay/ancient-pheno-pred/.

