## Supplementary Figures for "Natural selection acting on complex traits hampers the predictive accuracy of polygenic scores in ancient samples"

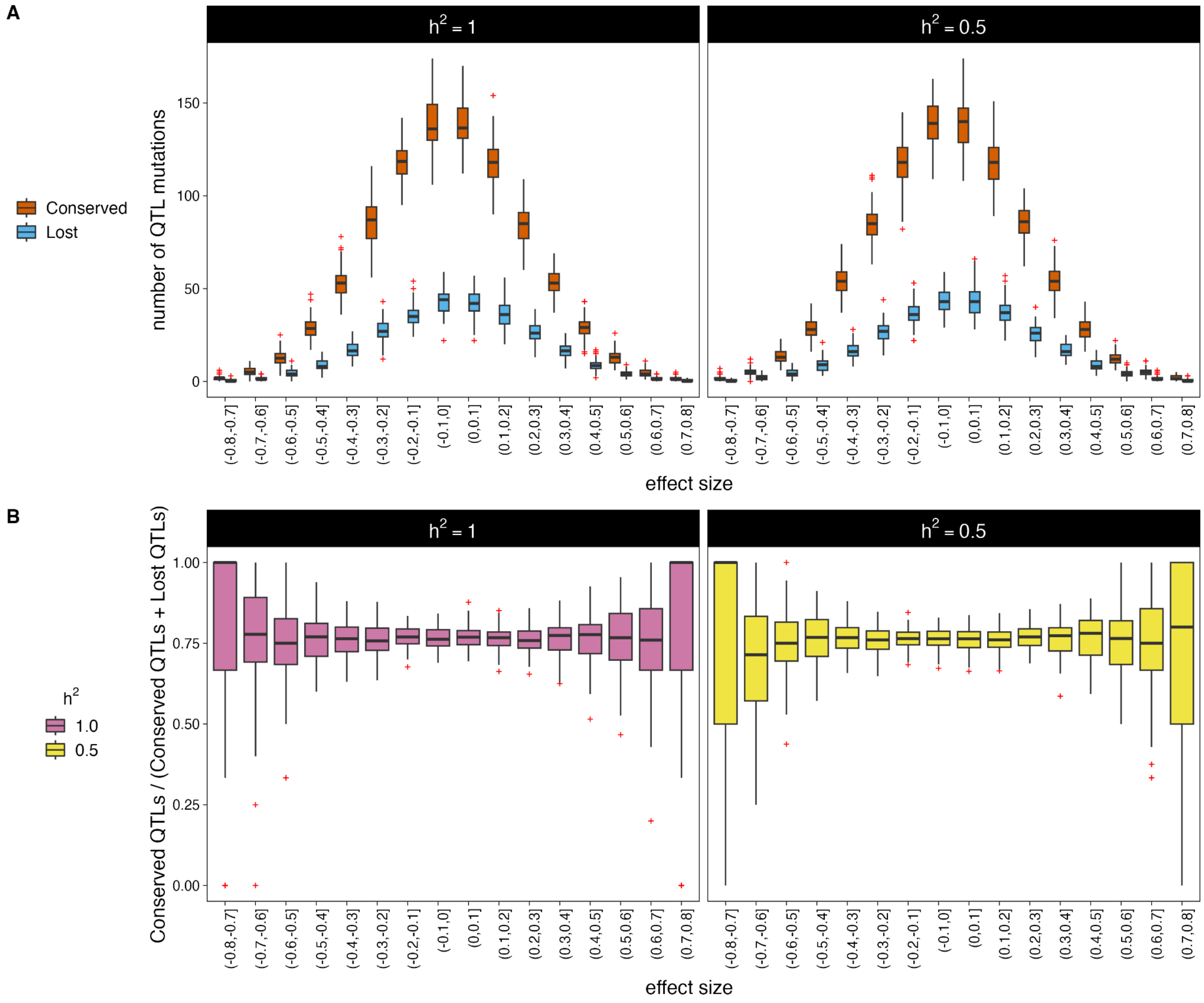


**Figure S1. A)** Effect sizes (X-axis) of the number of QTL mutations (Y-axis) that are conserved (orange) and lost (blue) between the earliest sampling time and the present-day sampling time ($\tau = 400$ and $0$ generations ago, respectively) in a sample of $100$ individuals taken at time $\tau = 400$ and another sample of $100$ individuals taken at time $\tau = 0$. **B)** The number of conserved QTL mutations divided by the total number of QTL mutations (conserved QTLs plus lost QTLs) (X-axis) per effect size bin (X-axis) between the earliest sampling time and the present-day sampling time ($\tau = 400$ and $0$ generations ago, respectively). Results are shown for 100 simulation replicates with heritability values of $h^2 = 1.0$ (left) and $h^2 = 0.5$ (right). Red crosses represent outliers.

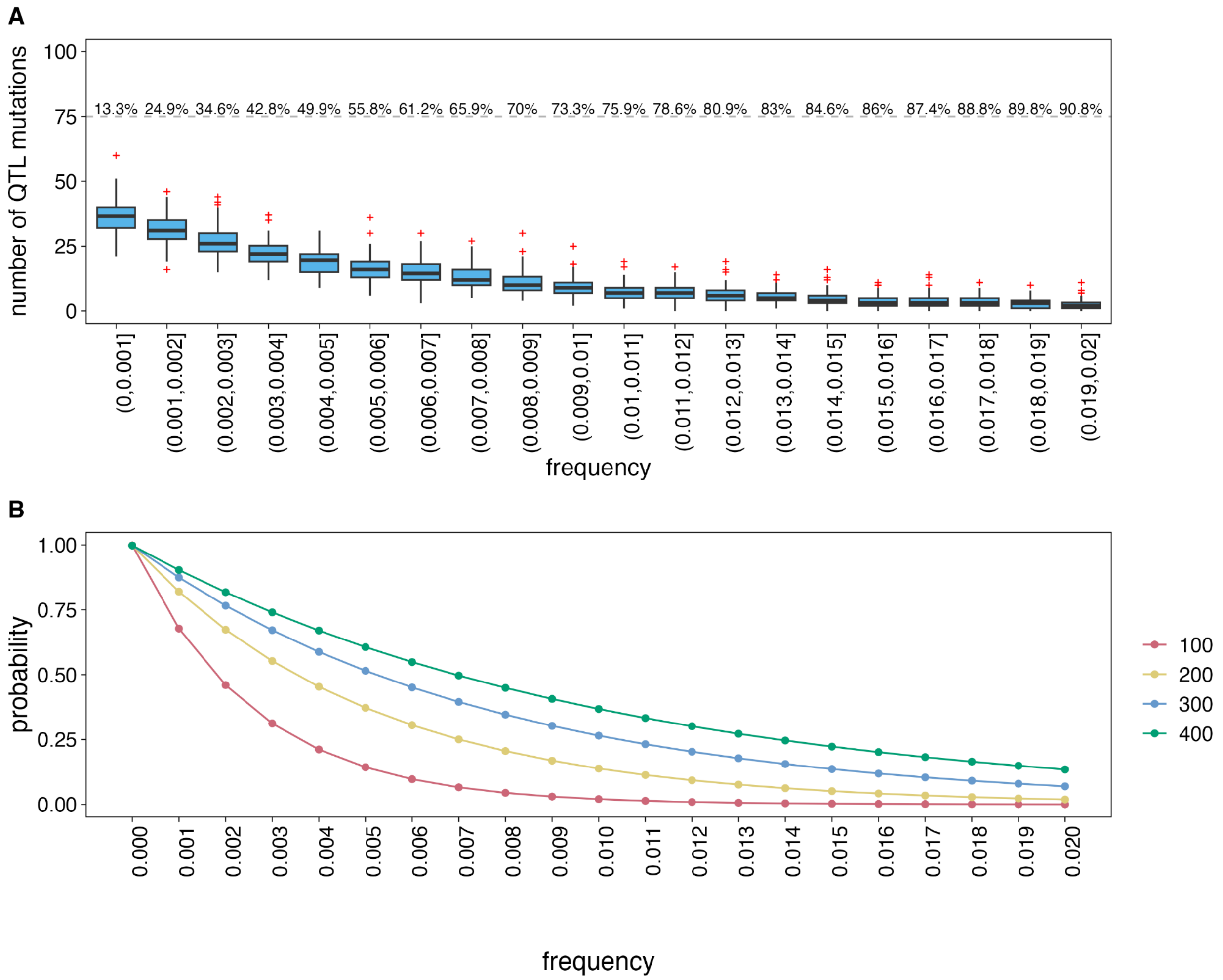


**Figure S2. A)** The population allele frequency of ~$90\%$ of simulated mutations that are lost between the earliest sampling time and the present-day sampling time ($\tau = 400$ and $0$ generations ago, respectively). The percentages above the gray dashed line represent the cumulative percentage sum (starting from the lower frequency) of the number of lost QTLs in each bin relative to the total number of lost QTLs. **B)** The probability of transitioning from an initial frequency, $f$, (x-axis) to a frequency of $0$ in $100, 200, 300$ and $400$ generations in pink, yellow, blue and green, respectively.

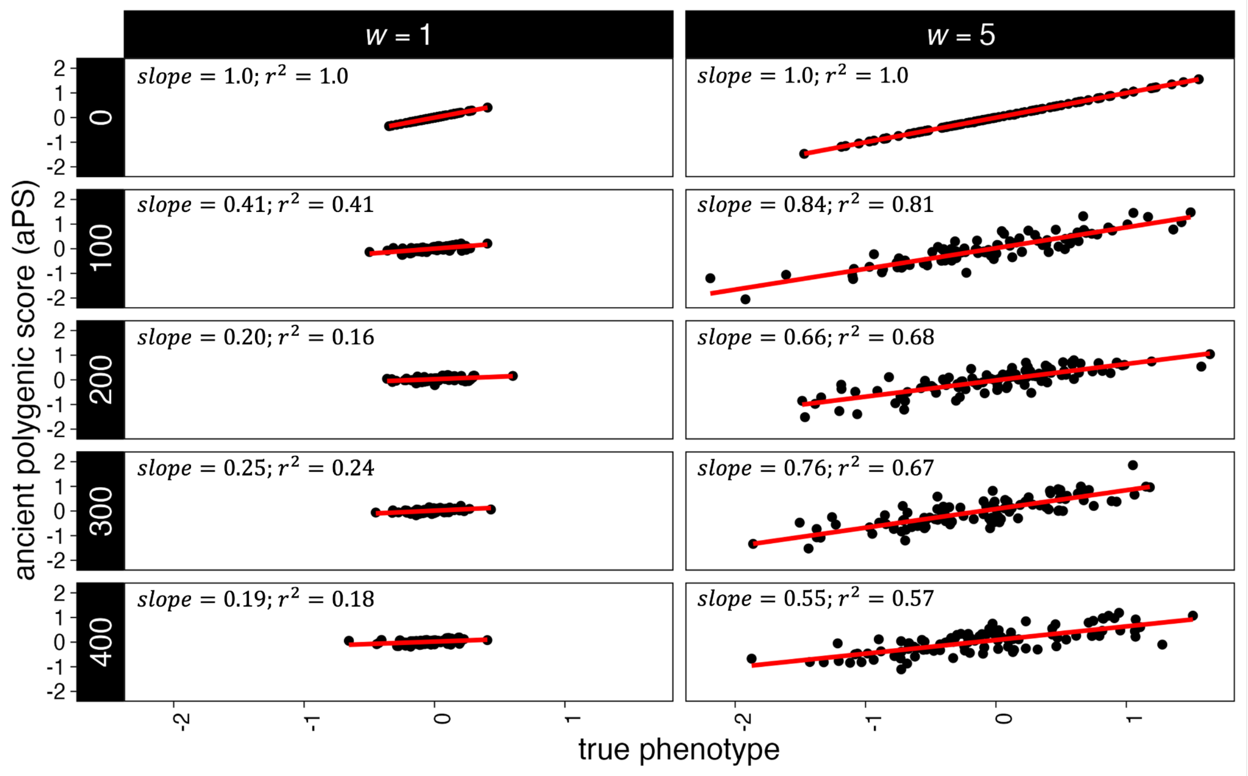


**Figure S3.** The true phenotype (X-axis) and the predicted ancient polygenic score (Y-axis) of a sample of $100$ individuals taken $0, 100, 200, 300$ and $400$ generations (rows) before the present at two different strengths of stabilizing selection representing both strong ($w = 1$, left), and weak ($w = 5$, right) selection acting on the simulated trait. The results shown in panels come from a single simulation replicate.

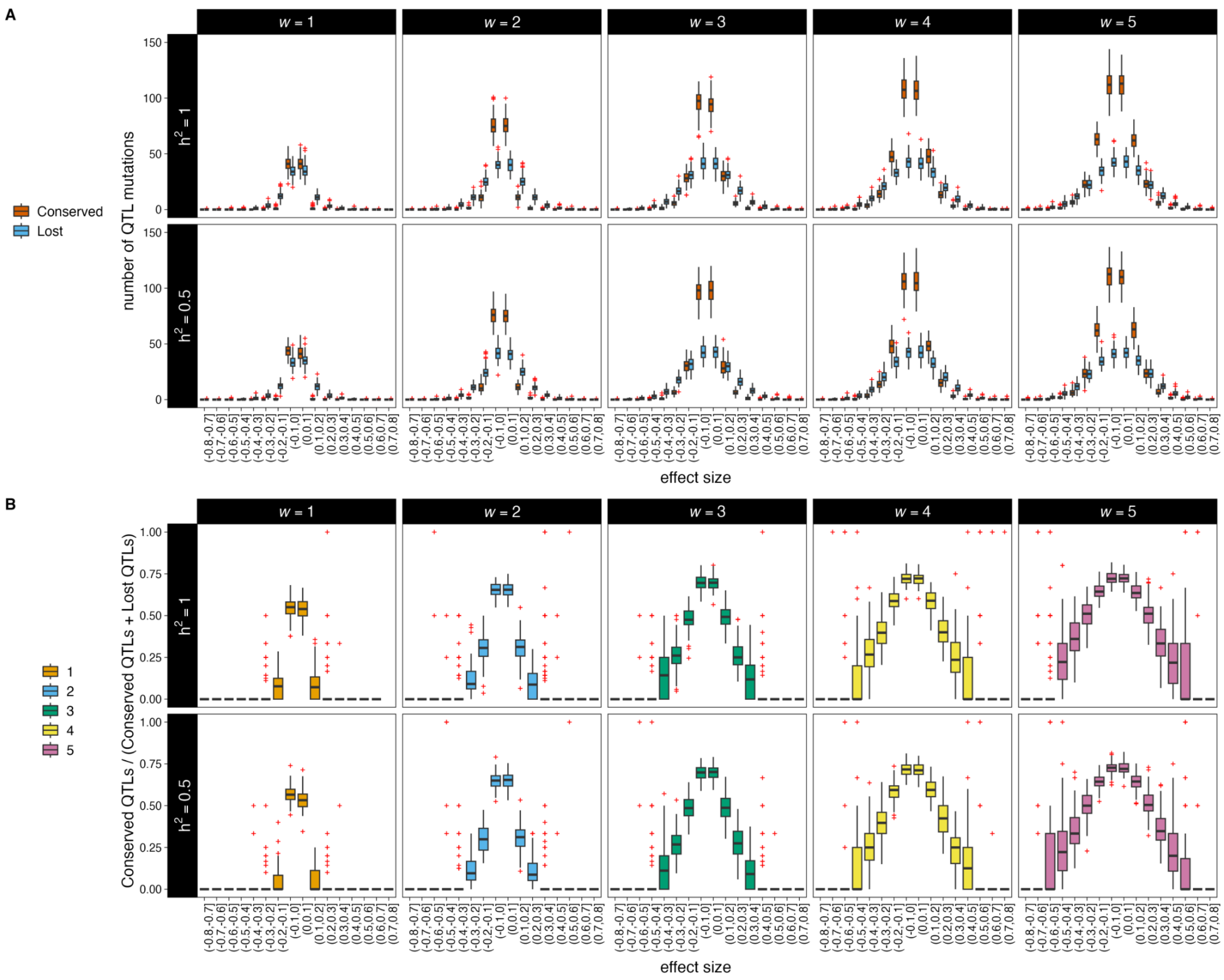


**Figure S4. A)** Effect sizes (X-axis) of the number of QTL mutations (Y-axis) that are conserved (orange) and lost (blue) between the earliest sampling time and the present-day sampling time ($\tau = 400$ and $0$ generations ago, respectively) in a sample of $100$ individuals taken at time $\tau = 400$ and another sample of $100$ individuals taken at time $\tau = 0$. **B)** The number of conserved QTL mutations divided by the total number of QTL mutations (conserved QTLs plus lost QTLs) (X-axis) per effect size bin (X-axis) between the earliest sampling time and the present-day sampling time ($\tau = 400$ and $0$ generations ago, respectively). Results are shown for $100$ replicates at heritability values of $h^2 = 1.0$ (top) and $h^2 = 0.5$ (bottom) at the five different strengths of selection ranging from strong to weak selection, $w = \{1, 2, 3, 4, 5\}$. Red crosses represent outliers.

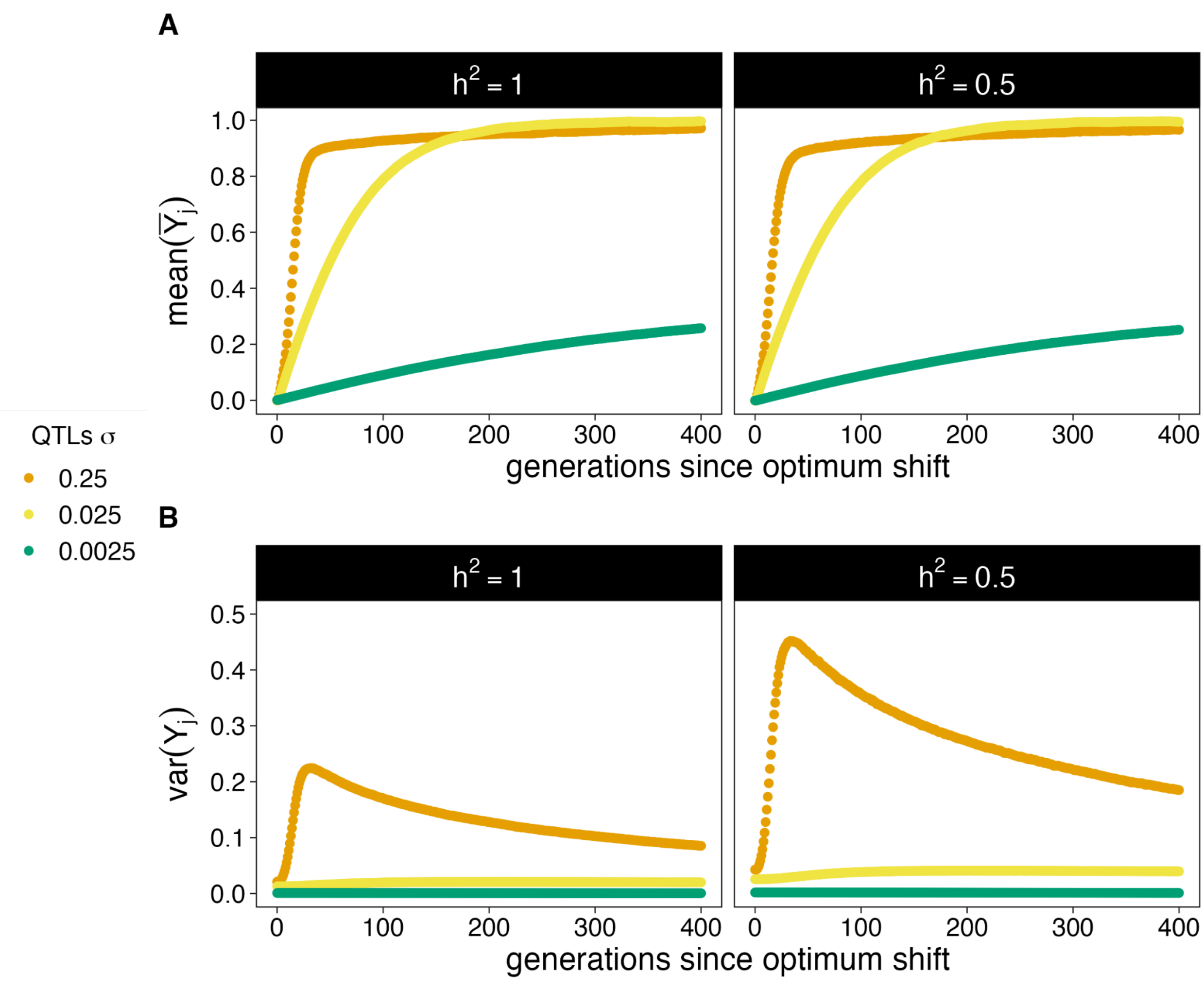


**Figure S5. Average phenotype mean and average phenotypic variance through time.** We model a trait with a heritability value of $h^2 = 1.0$ (left) and $h^2 = 0.5$ (right) evolving under stabilizing selection with a parameter $w = 1$. We imposed an optimum shift from $Y_O = 0$ to $Y_O{}' = 1$ in $400$ generations after a burn-in period and tested three different standard deviations of the QTL effect sizes distribution, $QTL \sim N(\mu = 0, \sigma = \{0.25, 0.025, 0.0025\})$, (in orange, yellow and green, respectively). The X-axis shows the number of generations since the optimum shift was imposed. The Y-axis shows the average phenotype mean in **A)** and the variance of phenotypic values in **B)**. We computed each average metric over the $100$ replicates we ran for each parameter combination.

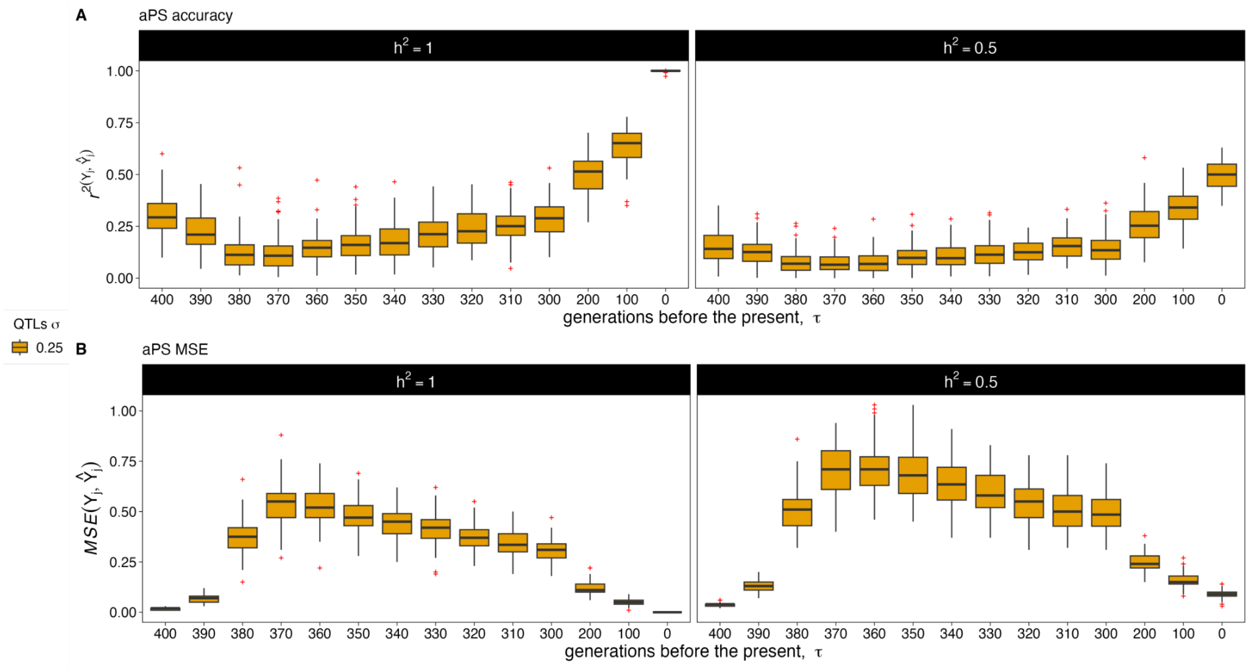


**Figure S6. Ancient polygenic scores (aPS) accuracy (*r*²) and Mean Squared Error (MSE) for a trait evolving under directional selection.** We model a trait with a heritability value of $h^2 = 1.0$ (left) and $h^2 = 0.5$ (right) evolving under directional selection with an optimum shift from $Y_O = 0$ to $Y_O{}' = 1$ 400 generations ago. Boxplots show the distribution of **A)** $r^2(Y, \hat{Y})$, and **B)** the Mean Squared Error (MSE), $MSE(Y, \hat{Y})$, between the true phenotypic values and their predicted ancient polygenic scores of a sample of $100$ individuals at different points in time, $\tau = 0, 100, 200, 300, 310, 320, 330, 340, 350, 360, 370, 380, 390$ and $400$ generations before the present, over 100 simulation replicates. Red crosses represent outliers.

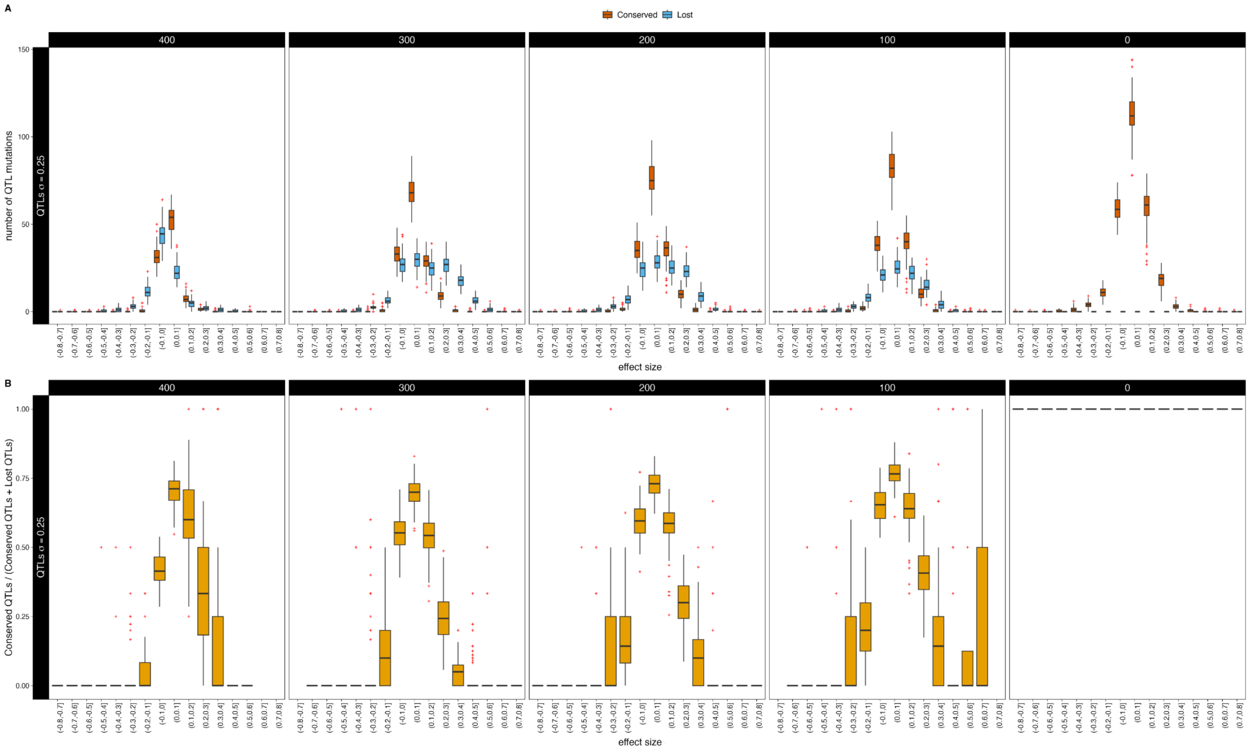


**Figure S7. Conserved and Lost QTL mutation patterns when simulating a trait evolving under directional selection with a heritability value $h^2 = 1.0$ and a QTL standard deviation of effect sizes $\sigma = 0.25$. A)** Effect sizes (X-axis) of the number of QTL mutations (Y-axis) that are conserved (orange) and lost (blue) between the different ancient sampling times and the present-day time. **B)** The number of conserved QTL mutations divided by the total number of QTL mutations (conserved QTLs plus lost QTLs) (Y-axis) per effect size bin (X-axis) between the different ancient sampling times and the present-day time. Each column uses $100$ individuals taken from a different sampling time, $\tau = 400, 300, 200, 100, 0$ generations ago, respectively. Results are shown for $100$ replicates. Red crosses represent outliers.

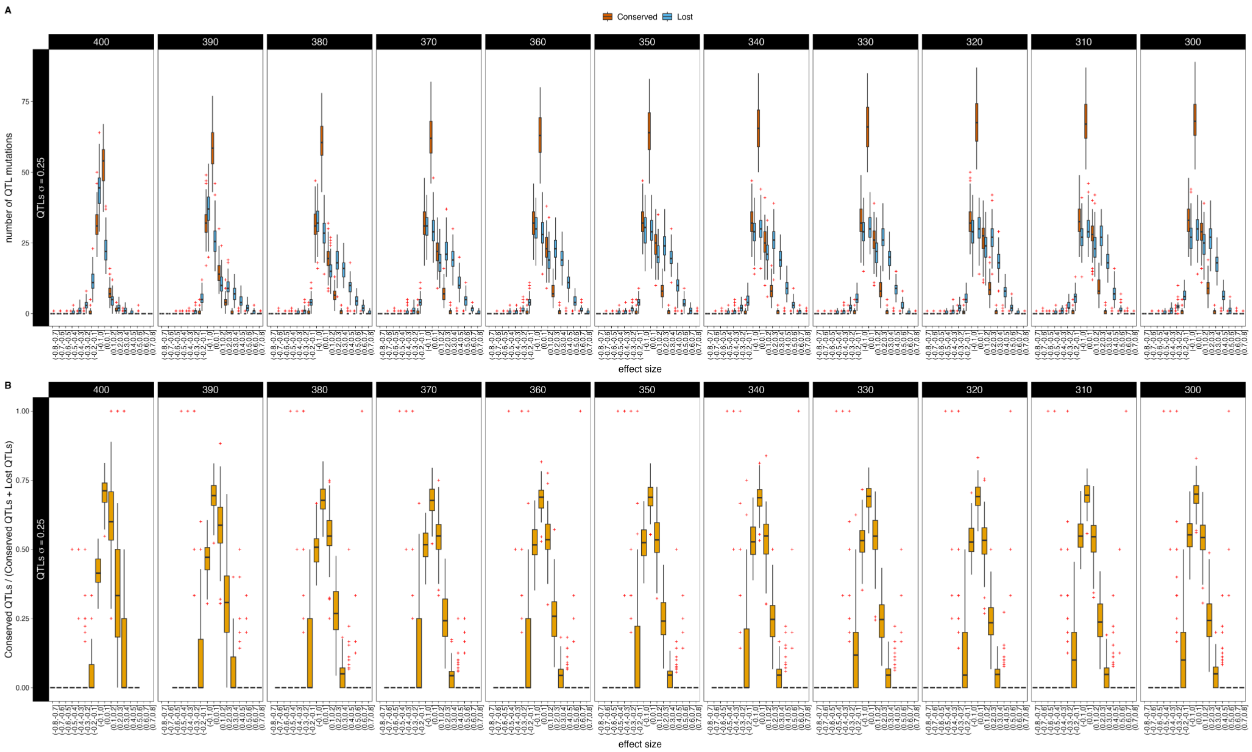

**Figure S8. Conserved and Lost QTL mutation patterns when simulating a trait evolving under directional selection with a heritability value with a $h^2 = 1.0$ and a QTL standard deviation of effect sizes $\sigma = 0.25$.** Generation times range from $\tau = 400$ to $\tau = 300$. **A)** Effect sizes (X-axis) of the number of QTL mutations (Y-axis) that are conserved (orange) and lost (blue) between the different ancient sampling times and the present-day time. **B)** The number of conserved QTL mutations divided by the total number of QTL mutations (conserved QTLs plus lost QTLs) (Y-axis) per effect size bin (X-axis) between the different ancient sampling times and the present-day time. Each column uses $100$ individuals taken from a different sampling time, $\tau = 400, 390, 380, 370, 360, 350, 340, 330, 320, 310, 300$ generations ago, respectively. Results are shown for $100$ replicates. Red crosses represent outliers.

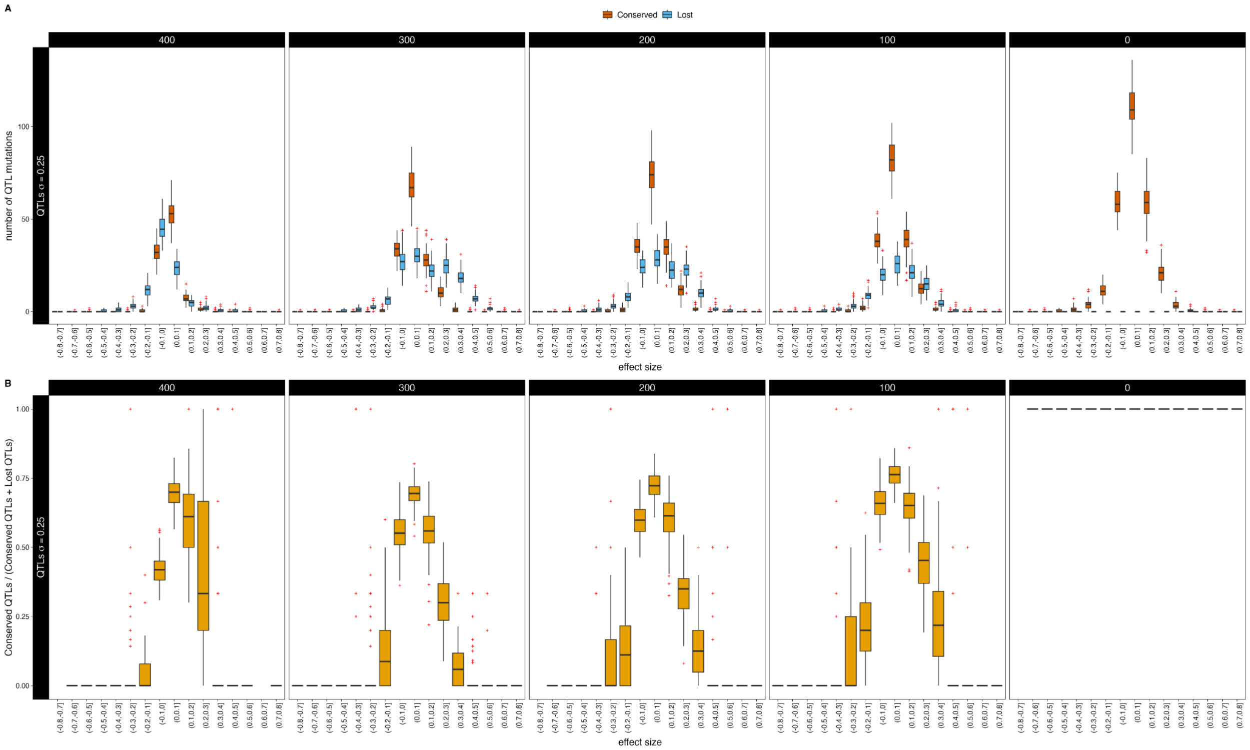


**Figure S9. Conserved and Lost QTLs patterns when simulating a trait evolving under directional selection with a heritability value of $h^2 = 0.5$ and a QTL standard deviation of effect sizes $\sigma = 0.25$. A)** Effect sizes (X-axis) of the number of QTL mutations (Y-axis) that are conserved (orange) and lost (blue) between the different ancient sampling times and the present-day time. **B)** The number of conserved QTL mutations divided by the total number of QTL mutations (conserved QTLs plus lost QTLs) (Y-axis) per effect size bin (X-axis) between the different ancient sampling times and the present-day time. Each column uses $100$ individuals taken from a different sampling time, $\tau = 400, 300, 200, 100, 0$ generations ago, respectively. Results are shown for $100$ replicates. Red crosses represent outliers.

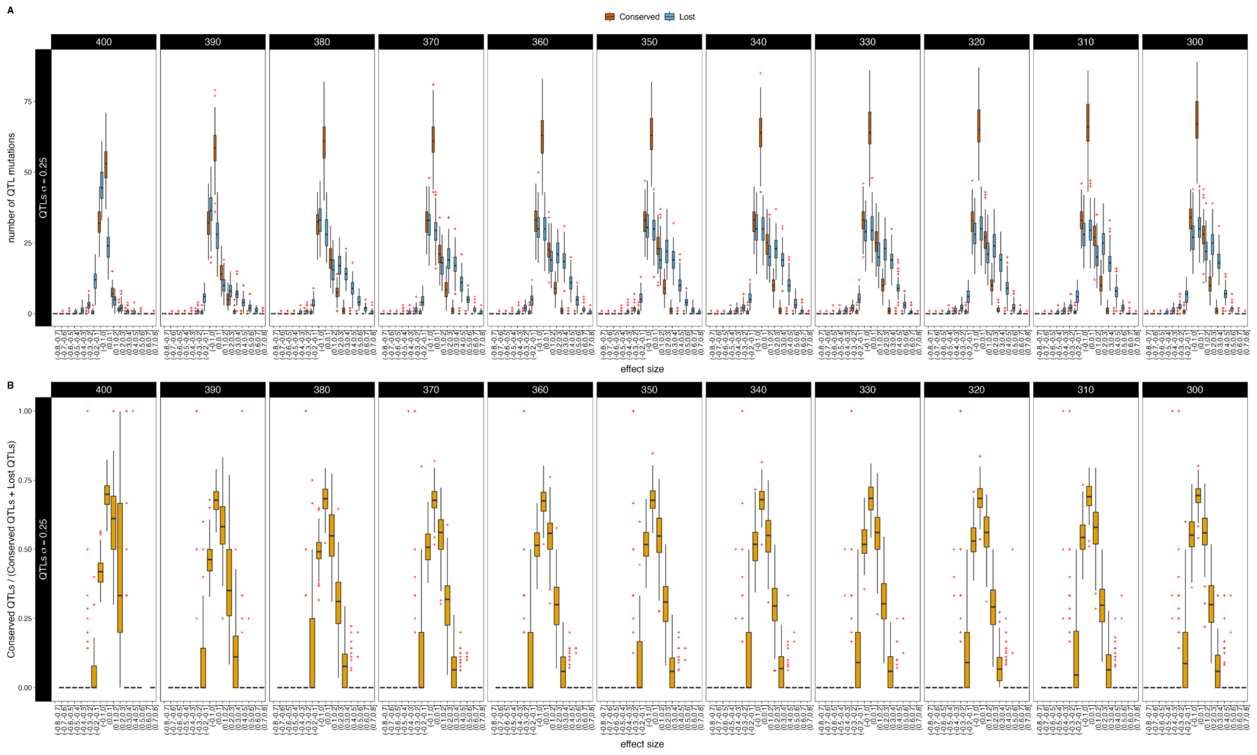


**Figure S10. Conserved and Lost QTLs patterns when simulating a trait evolving under directional selection with a heritability value of $h^2 = 0.5$ and a QTL standard deviation of effect sizes is $\sigma = 0.25$.** Generation times range from $\tau = 400$ to $\tau = 300$. **A)** Effect sizes (X-axis) of the number of QTL mutations (Y-axis) that are conserved (orange) and lost (blue) between the different ancient sampling times and the present-day time. **B)** The number of conserved QTL mutations divided by the total number of QTL mutations (conserved QTLs plus lost QTLs) (Y-axis) per effect size bin (X-axis) between the different ancient sampling times and the present-day time. Each column uses $100$ individuals taken from a different sampling time, $\tau = 400, 390, 380, 370, 360, 350, 340, 330, 320, 310, 300$ generations ago, respectively. Results are shown for $100$ replicates. Red crosses represent outliers.

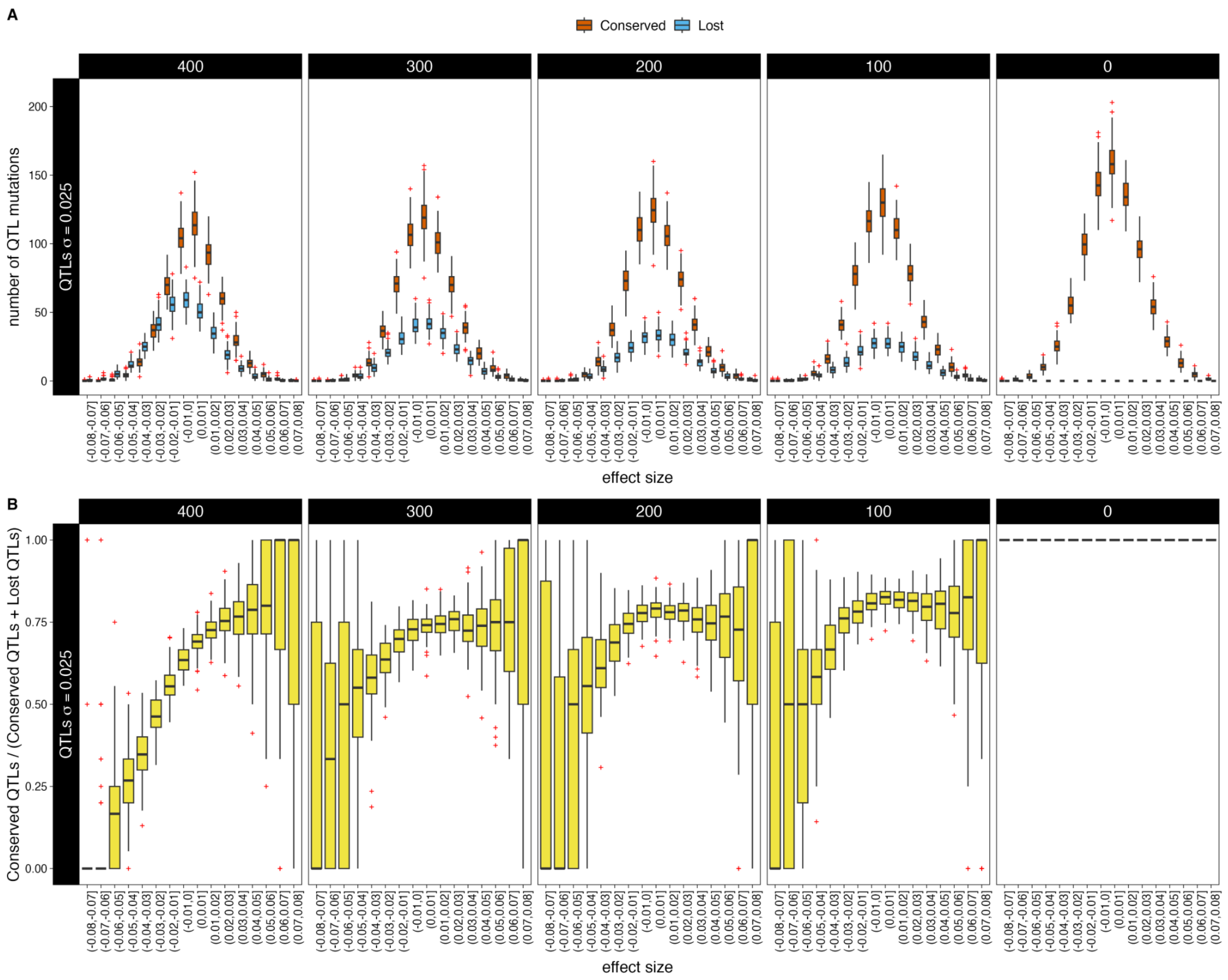


**Figure S11. Conserved and Lost QTLs patterns when simulating a trait evolving under directional selection with a heritability value of $h^2 = 1.0$ and a QTL standard deviation of effect sizes $\sigma = 0.025$. A)** Effect sizes (X-axis) of the number of QTL mutations (Y-axis) that are conserved (orange) and lost (blue) between the different ancient sampling times and the present-day time. **B)** The number of conserved QTL mutations divided by the total number of QTL mutations (conserved QTLs plus lost QTLs) (Y-axis) per effect size bin (X-axis) between the different ancient sampling times and the present-day time. Each column uses $100$ individuals taken from a different sampling time, $\tau = 400, 300, 200, 100, 0$ generations ago, respectively. Results are shown for $100$ replicates. Red crosses represent outliers.

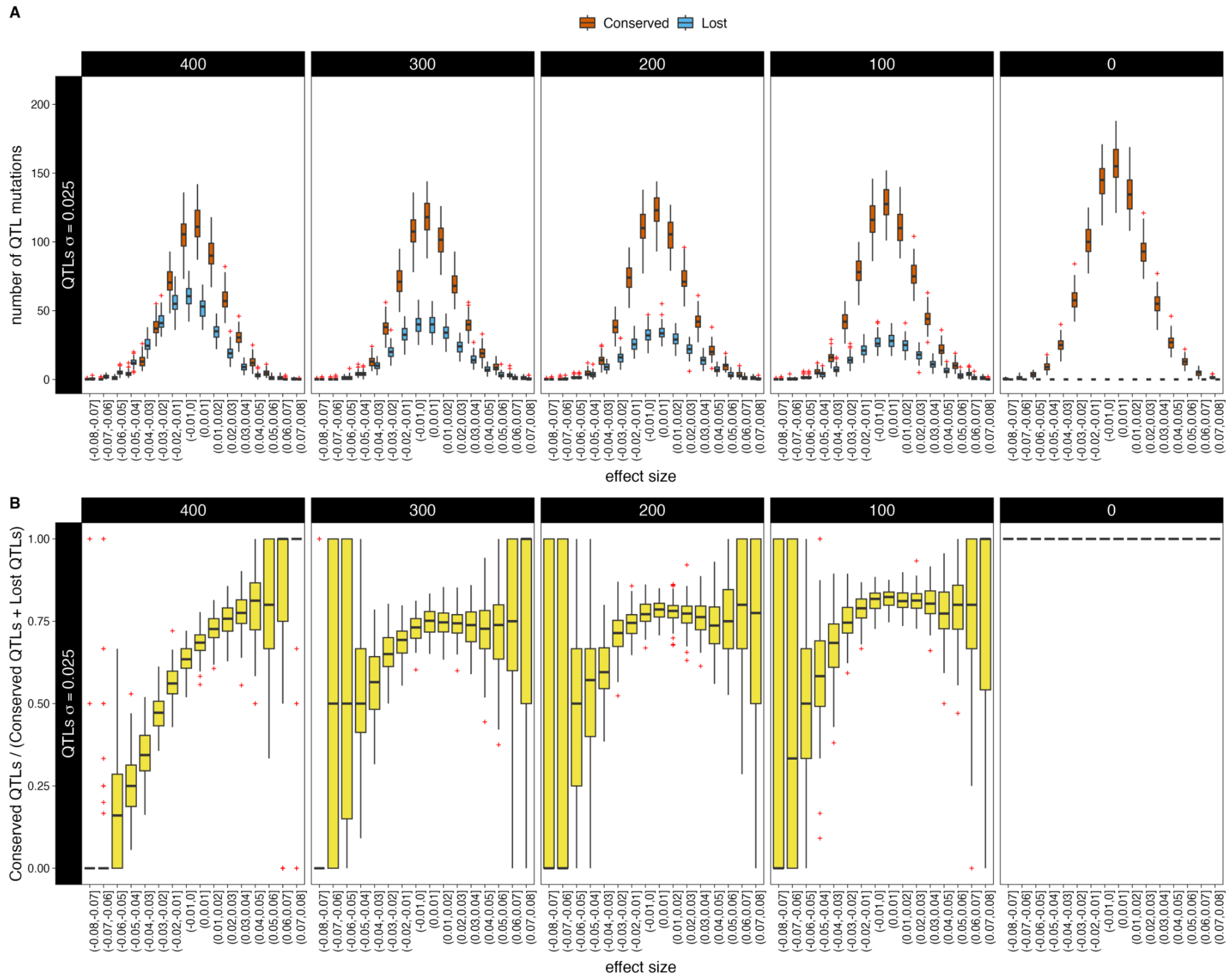


**Figure S12. Conserved and Lost QTLs patterns when simulating a trait evolving under directional selection with a heritability value of $h^2 = 0.5$ and a QTL standard deviation of effect sizes $\sigma = 0.025$. A)** Effect sizes (X-axis) of the number of QTL mutations (Y-axis) that are conserved (orange) and lost (blue) between the different ancient sampling times and the present-day time. **B)** The number of conserved QTL mutations divided by the total number of QTL mutations (conserved QTLs plus lost QTLs) (Y-axis) per effect size bin (X-axis) between the different ancient sampling times and the present-day time. Each column uses $100$ individuals taken from a different sampling time, $\tau = 400, 300, 200, 100, 0$ generations ago, respectively. Results are shown for $100$ replicates. Red crosses represent outliers.

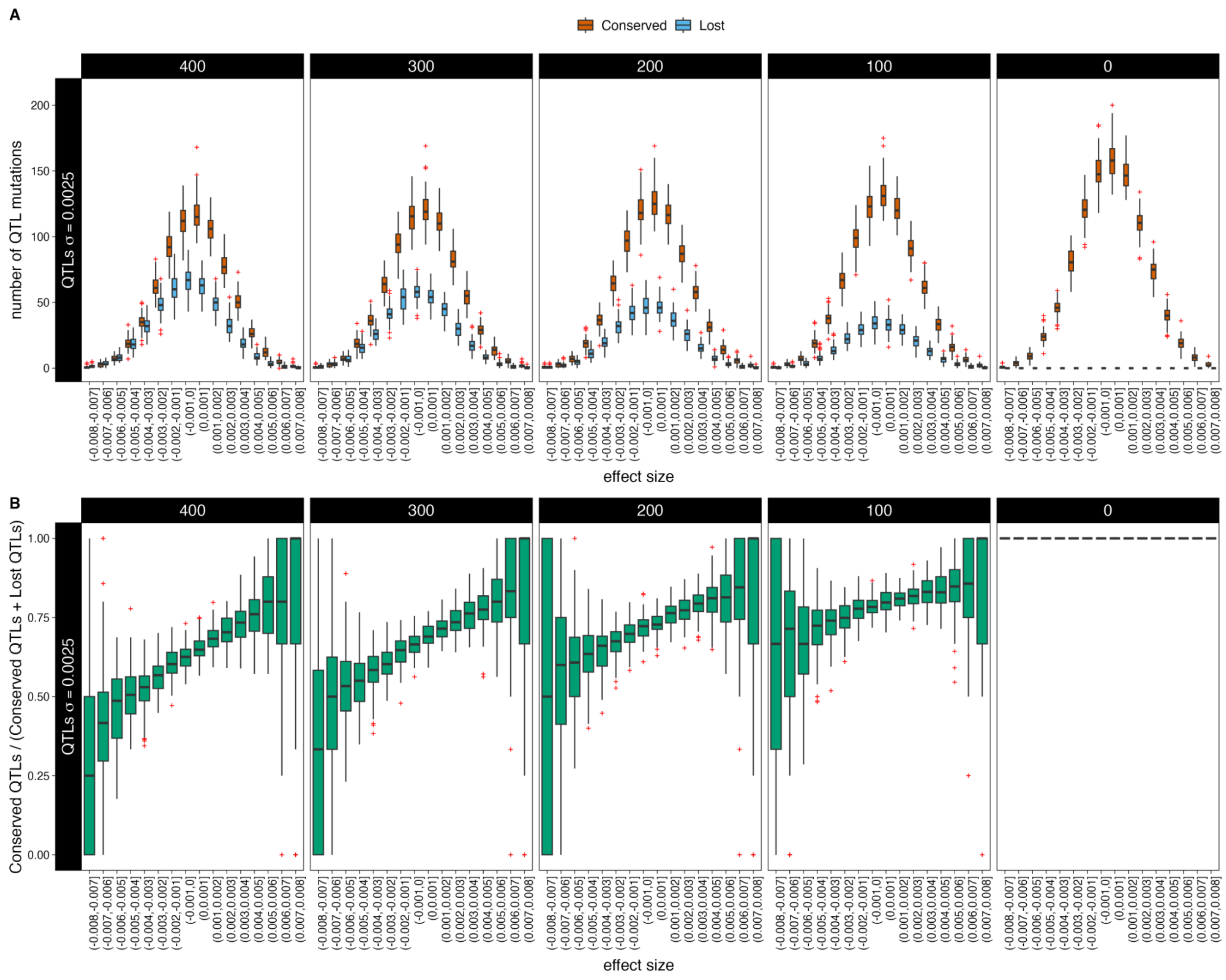


**Figure S13. Conserved and Lost QTLs patterns when simulating a trait evolving under directional selection with a heritability value of $h^2 = 1.0$ and a QTL standard deviation of effect sizes $\sigma = 0.0025$. A)** Effect sizes (X-axis) of the number of QTL mutations (Y-axis) that are conserved (orange) and lost (blue) between the different ancient sampling times and the present-day time. **B)** The number of conserved QTL mutations divided by the total number of QTL mutations (conserved QTLs plus lost QTLs) (Y-axis) per effect size bin (X-axis) between the different ancient sampling times and the present-day time. Each column uses $100$ individuals taken from a different sampling time, $\tau = 400, 300, 200, 100, 0$ generations ago, respectively. Results are shown for $100$ replicates. Red crosses represent outliers.

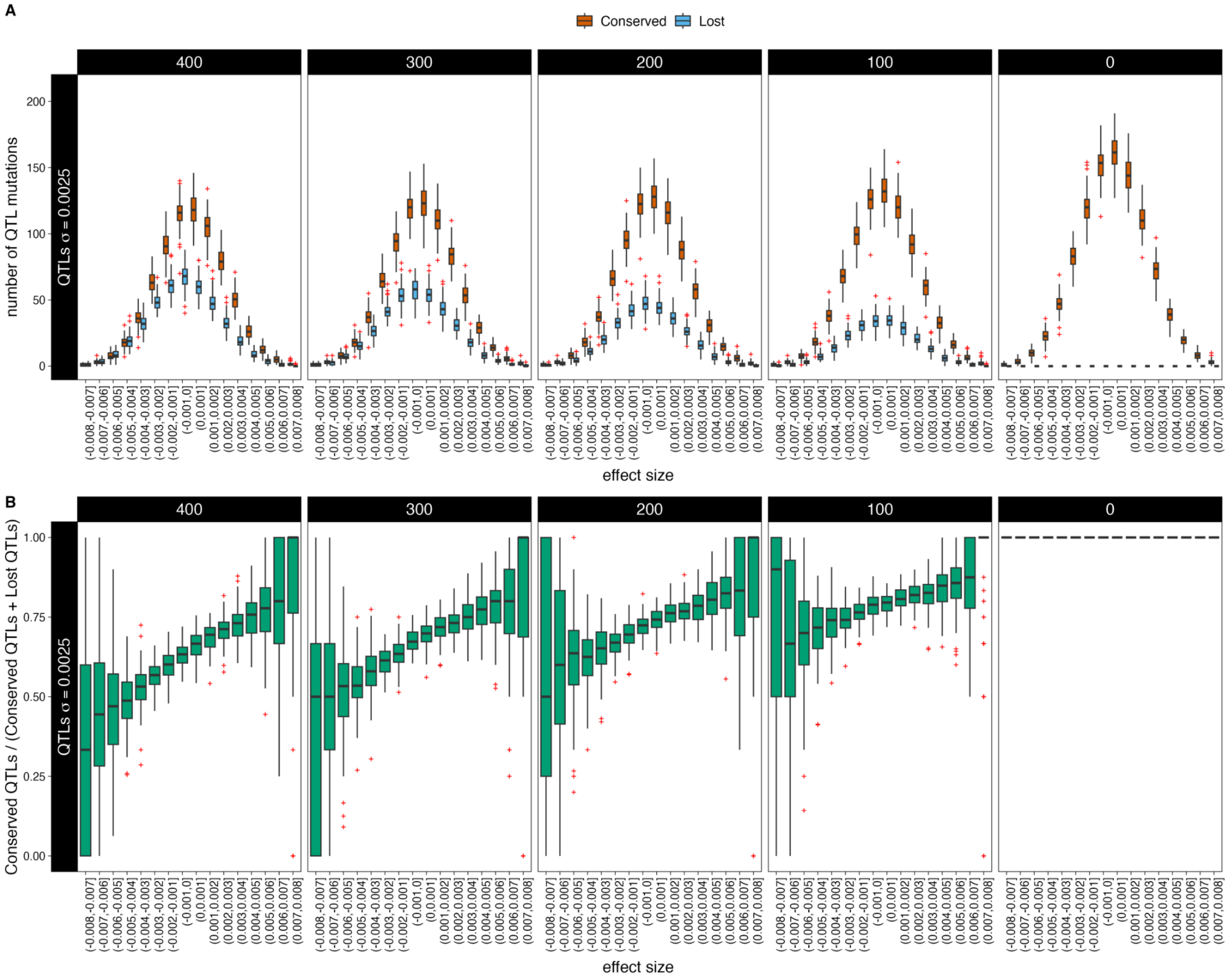


**Figure S14. Conserved and Lost QTLs patterns when simulating a trait evolving under directional selection with a heritability value of $h^2 = 0.5$ and a QTL standard deviation of effect sizes $\sigma = 0.0025$. A)** Effect sizes (X-axis) of the number of QTL mutations (Y-axis) that are conserved (orange) and lost (blue) between the different ancient sampling times and the present-day time. **B)** The number of conserved QTL mutations divided by the total number of QTL mutations (conserved QTLs plus lost QTLs) (Y-axis) per effect size bin (X-axis) between the different ancient sampling times and the present-day time. Each column uses $100$ individuals taken from a different sampling time, $\tau = 400, 300, 200, 100, 0$ generations ago, respectively. Results are shown for $100$ replicates. Red crosses represent outliers.
